# Oligopeptide (FK)_4_ facilitates the determination of the absolute stereochemistry of chiral organic compounds *via* NMR Spectroscopy

**DOI:** 10.1101/2024.06.17.599446

**Authors:** Mingjun Zhu, Yun Peng, Kaihua Zhang, Jiening Wang, Gangjin Yu, Guan Wang, Shan Wu, Zhou Gong, Xu Zhang, Lichun He, Maili Liu

## Abstract

Molecular chirality plays a crucial role in the fields of chemistry and biology. Various nuclear magnetic resonance (NMR) methods have been employed for assignment of stereochemical configurations. Alignment media for measurements of anisotropic parameters contribute to configuration elucidations, while chiral auxiliaries assist in enantiomer discrimination and chiral recognition. However, each method comes with limitations, and the assignment of absolute configuration by NMR remains challenging. In this study, we explored a combined approach for absolute configuration elucidation with the multifaceted application of the oligopeptide (FK)_4_, which possesses dual functionalities as an alignment medium for anisotropic NMR measurements and a chiral differentiating agent. As an alignment medium, (FK)_4_ facilitates the analysis of stereochemical features through residual dipolar couplings (RDC), aiding in the structural elucidation of stereoisomers. Moreover, (FK)_4_ exhibits unique chiral differentiating properties, interacting distinctly with enantiomers and resulting in observable bifurcation of chemical shift signals, in ^13^C, ^1^H and ^19^F spectra. Using isoleucine and xylose as model compounds, we demonstrate how the dual functionality of (FK)_4_ enables the identification of stereoisomeric structures *via* RDC parameters and the assignment of enantiomeric configurations through measurements of t2 relaxation time coupled with theoretical simulations. These findings highlight the potential of versatile chiral media as (FK)_4_ for approaching the elucidation of absolute configurations of organic molecules *via* combined NMR spectroscopy approaches, with implications for structural characterization and enantiomer discrimination in chemical applications.

## Introduction

Molecular chirality is pivotal in chemical^1-3^, physical^4^, pharmaceutical^5^, and biology^6^, with biological homochirality, such as L-amino acid and D-sugars, fundamental to life on Earth. X-ray crystallography is a primary method for assignment of absolute configurations, though obtaining high-quality crystals, especially for light atoms, poses challenges^7^. Nuclear magnetic resonance (NMR) spectroscopy offers an alternative approach for structural elucidation and chiral recognition, especially for non-volatile or thermally unstable compounds^8, 9^. Alignment media such as lyotropic liquid crystals aid in measurements of anisotropic NMR parameters, specifically residual dipolar coupling (RDC) and residual chemical shift anisotropy (RCSA). Complemented by density functional theory (DFT), unequivocal configuration and conformation of organic molecules can be elucidated^10-17^. However, distinguishing between mirror-imaged enantiomers remains challenging due to the complexity of theoretical calculations predicting the miniscule change in stereoisomeric conformations and respected chemical shifts of the analyte molecules while taking into account the alignment medium^18, 19^. On the other hand, converting enantiomers to diastereomers using a chiral auxiliary such as a chiral derivatization agent (CDA)^20-22^ or chiral solvating agent (CSA)^8, 22-25^ facilitates successful chiral recognition. This feature of the chiral differentiating agent could potentially enhance the elucidation of absolute configuration when combined with the characteristic of alignment media. Current chiral auxiliaries mainly consist of small molecules that freely diffuse in the solution environment^8, 20-25^, while reported alignment media usually demonstrate equivalent interactions with a pair of enantiomers^26, 27^. The lack of suitable agents limits current applications in conjunction.

In our prior research, we utilized the oligopeptide (FK)_4_ as an alignment medium for anisotropic measurements to determine molecular configurations^28, 29^. Here, we further investigated the intermolecular interactions between (FK)_4_ and analytes including amino acids, sugars, organic acids and commercially available drugs. Due to different hydrogen bonding patterns within (FK)_4_, this oligopeptide can serve as a chiral differentiating alignment media for elucidating absolute configuration of aminos acids and sugars. Furthermore, we proposed a combined NMR method that first identifies stereoisomeric structures of sample analytes using RDC measurements, and then assigns the enantiomeric configurations through t2 relaxation time analysis coupled with DFT calculations. Additionally, we demonstrated that (FK)_4_ can effectively recognize enantiomers of various biological active small molecules and representative commercially available drugs with accurate determination of enantiopurity. Leveraging the properties of lyotropic liquid crystal alignment media in anisotropic solvent environments and as a chiral solvating agent for intermolecular interactions with analytes, (FK)_4_ emerges as a novel chiral differentiation agent poised for elucidating the absolute configuration of small molecules and extensive applications in chiral recognition.

## Results and Discussion

### Interactions between the self-assembled liquid crystal chiral differentiating agent (FK)_4_ and amino acid analytes

The oligopeptide (FK)_4_ consists of 4 repeating unit of two alternating amino acids, phenylalanine (F) and lysine (K) (Figure 1A left). The aromatic phenyl group in phenylalanine imparts hydrophobic properties, facilitating π-π stacking interactions. Conversely, the amine moiety in lysine confers hydrophilic characteristics and is prone to protonation, resulting in a positive charge under acidic conditions. The binary patterning of polar and nonpolar residues in (FK)_4_ derived the formation of fibrils resembling amyloid, which further self-assembled into antiparallel β-sheet structures (Figure 1A)^30-35^. Transmission electron microscopy (TEM) and cryo-electron microscopy imaging of self-assembled (FK)_4_ revealed thin rigid fibrils, suggesting its potential to segment the solution environment with directional alignment (Figure 1B left and Figure S1). The presence of a meridional reflection at approximately 4.7 Å, observed in the Fourier transform (FT) of the TEM images, confirmed the formation of a β-sheet secondary structure post self-assembly (Figure 1B right)^31^. The inherent chirality of (FK)_4_, with both phenylalanine and lysine residues in their natural Lenantiomer configuration, resembled characteristics of previously reported CSAs^8, 22-25^, indicating its potential capacity for chirality recognition.

**Figure 1.**
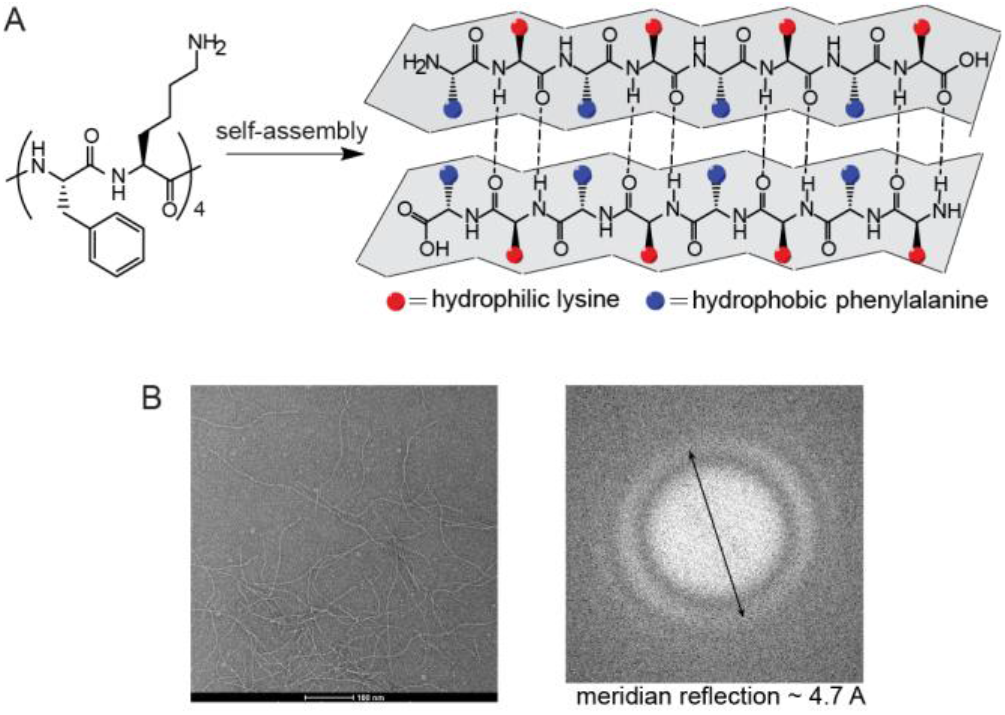
Characterization structure and secondary structure of oligopeptide (FK)_4_. B. TEM images of self-assembled oligopeptide (FK)_4_.

Towards this hypothesis, we first explored investigated enantiomer recognition in the presence of (FK)_4_. Enantiomers of both valine and isoleucine were distinguished using proton-decoupled ^13^C NMR, yielding distinct results (Figure 2A and 2B). Multiple carbons situated in diverse chemical environments within isoleucine displayed divergent resonances (Figure 2B), while valine exhibited a single signal at the carboxyl carbon under identical experimental conditions (Figure 2A). These differences may stem from interactions between (FK)_4_ and the amino acids. Spin-spin (t2) relaxation times were measured to investigate molecular interactions in the system. T2 relaxation indicates the process by which the magnetization in the equatorial plane decays to its equilibrium value of zero and is influenced by the chemical environment surrounding the targeted nuclei^36-38^. Generally, longer t2 value corresponds to less dephasing of the nuclei spins, which aligns with fewer interactions around the atom^39, 40^. For valine, L-enantiomer exhibited a slightly longer C2 t2 of 0.53 s, compared to D-enantiomer of 0.49 s, suggesting weaker interaction with (FK)_4_ centered at C2 for L-valine. In contrast, L-isoleucine showed a shorter C2 t2 of 0.26 s than D-isoleucine of 0.31 s, indicating stronger interaction for the L-enantiomer around C2 (Figure 2C and 2D). Interestingly, isoleucine enantiomers displayed a more pronounced t2 difference than valine, likely due to the electron-donating properties of the hydrocarbon side chain. The longer hydrocarbon side chain in isoleucine results in a more electron-rich carboxyl carbon, facilitating stronger hydrogen bonding with (FK)_4_.

**Figure 2.**
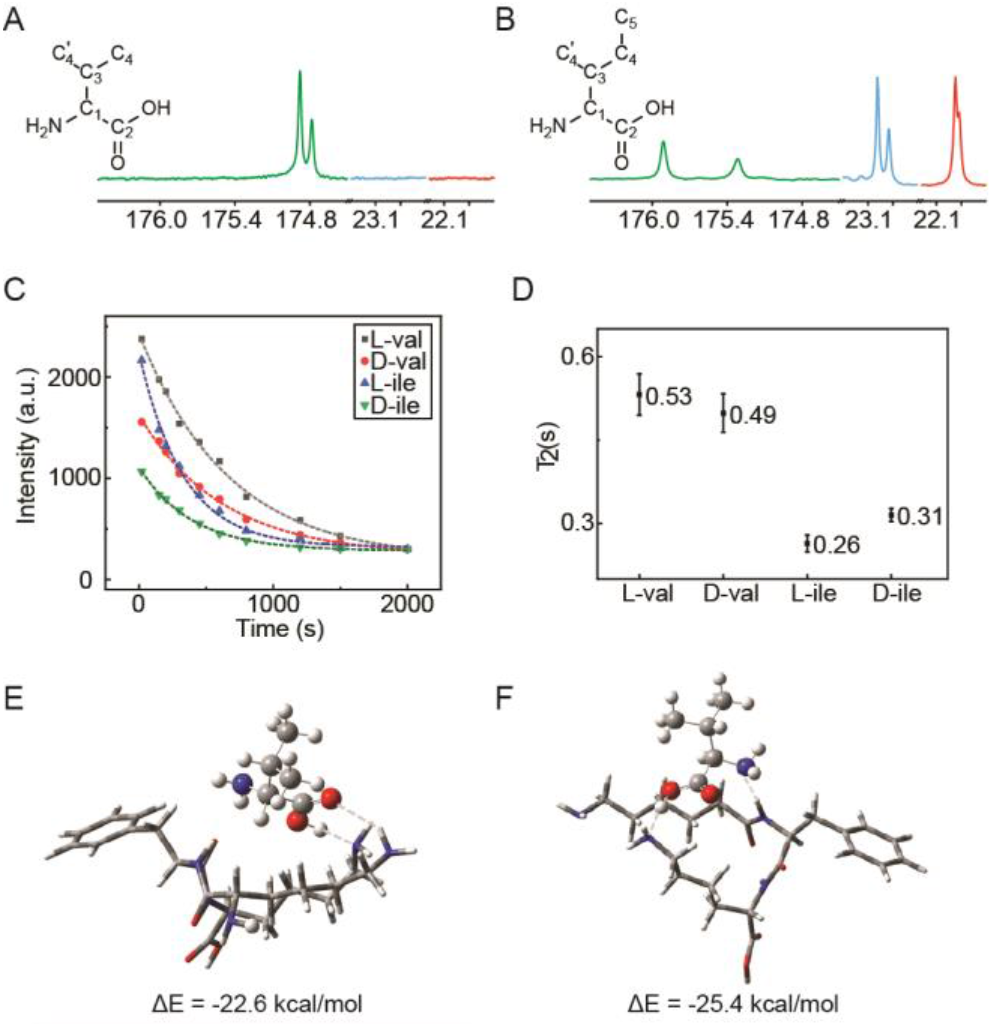
Interactions between amino acid analytes and (FK)_4_. A. Proton decoupled ^13^C spectrum of 65 mM valine enantiomers in presence of 140 mg/ml (FK)_4_. B. Proton decoupled ^13^C spectrum of 65 mM isoleucine enantiomers in the presence of 140 mg/ml (FK)_4_. C. T_2_ relaxation decay of 40 mM L-valine and 25 mM D-isoleucine enantiomers in the presence of 140 mg/ml (FK)_4_. D. T2 relaxation times extrapolated from respected NMR results. E. L-valine interacts with (FK)_2_ model simulated with DFT. F. D-valine interacts with (FK)_2_ model simulated with DFT. (FK)_2_ was represented with tube model, and valine was represented with ball-stick model.

Density functional theory (DFT) calculations were performed to further investigate the interaction between the amino acid analytes and the chiral differentiating agent. As (FK)_4_ are composed of repeated units, a simplified model (FK)_2_ featuring alternating phenylalanine and lysine units was used for simulation. Valine and isoleucine were introduced as sample molecules into the system. DFT calculations were carried out for multiple configurations post-interaction between the oligopeptide and the analyte, yielding stable structures corresponding to different enantioselectivities of valine (Figure 2E, 2F, and the corresponding structure files are listed in the SI 4.2). (FK)_2_ is expected to represent the interaction strength as the binding modes could be extended to (FK)_4_. In L-valine, stabilization occurred through interactions of the hydroxyl (OH) and oxo(O) of the analyte’s carboxyl group with the lysine chains, while the amine group (NH_2_) of the analyte was spatially displaced away from the oligopeptide and its associated lysine units (Figure 2E). Conversely, D-valine primarily involved the hydroxyl group (OH) of the carboxylate groups in the hydrogen bonding network, with the amine group (NH_2_) positioned for interaction with the oligopeptide (Figure 2F). The stronger interaction of D-valine with the oligopeptide was evidenced by a stability increase of more than 2.8 kcal/mol compared to L-valine, consistent with the observed t2 relaxation difference between the enantiomers, resulting in a discernible separation in the NMR signals. Similarly, in L-isoleucine, the amine group was distanced from the oligopeptide; whereas in D-isoleucine, the amine group was proximal to the oligopeptide and forms an additional hydrogen bond (Figure S5), resulting in a 3.1 kcal/mol less stable in electronic energy. This result aligned with t2 measurement, in which D-isoleucine exhibited 0.05 s longer relaxation time, indicating weaker interactions between it and the oligopeptide, compared to the L-enantiomer. This variation in interaction energy facilitated chiral separation of isoleucine enantiomers using (FK)_4_ as a chiral differentiating agent in NMR spectroscopy. The calculation of analyte to oligopeptide interaction energies compelled with the experimental result of t2 relaxation times, leading to the assignment of bifurcated chemical shifts to respective enantiomers.

Atoms are color coded respectively as red for O, blue for N, grey for C, and white for H; hydrogen bonding interactions are indicated by dashed lines.

### Structural elucidation of amino acid and sugar enantiomers

The consistency observed in t2 measurements and computational modeling of intermolecular interactions between analytes and (FK)_4_ suggested a potential application in elucidation of absolute configuration. Acting as an alignment media, (FK)_4_ facilitated anisotropy measurements like RDC, enabling the resolution of relative configuration among multiple stereoisomers. For instance, RDC measurements of L-isoleucine confirmed the SS stereo-configuration but not SR (Figure 3A). With its CSA-like properties, (FK)_4_ induced peak bifurcation in the 1D ^13^C NMR spectrum of isoleucine (Figure 2B). Combining t2 relaxation measurements and DFT calculations of hydrogen bonding modes to (FK)_4_, specific chemical shifts can be assigned to each enantiomer. For example, two signals around 175.5 ppm corresponded to the same carbon (C2) in isoleucine, with different t2 results indicating varying electron environments around the carboxyl carbon between L- and D-enantiomers. Longer t2 values suggested weaker interactions between the analyte centered around the measured carbon (FK)_4_. A molecular model mimicking isoleucine and (FK)_4_ can be established using DFT or similar theoretical methods to simulate the interactions between the analyte enantiomers and the oligopeptide. Due to chirality at the C1 carbon, the side chain of isoleucine interacted with (FK)_4_ *via* different sites, leading to distinct assembly modes and environments around C2. Theoretical calculations can guide the assignment of t2 measurements and ^13^C chemical shifts to corresponding enantiomers, elucidating the absolute configuration of L-isoleucine distinct from its D-enantiomer.

**Figure 3.**
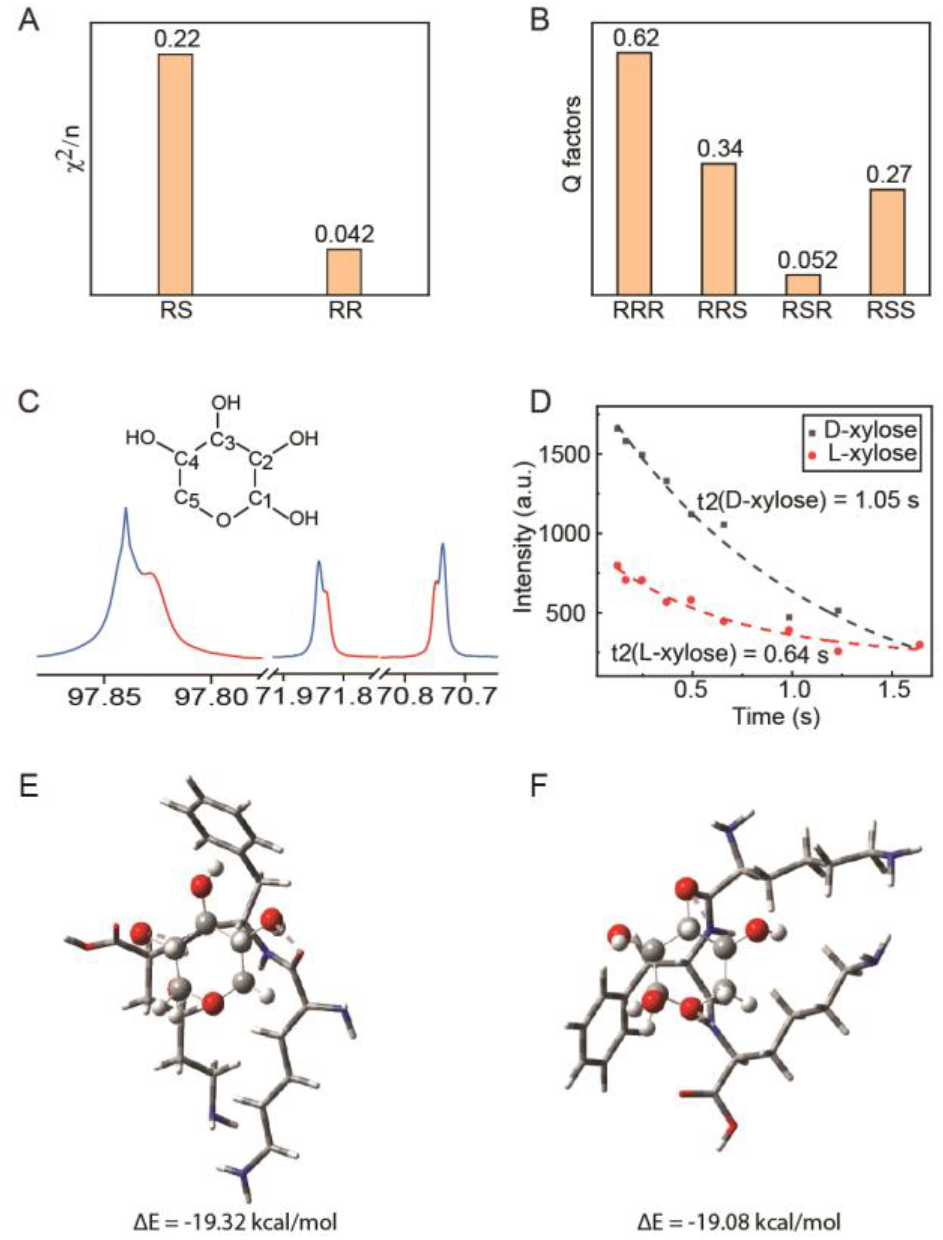
Structure elucidation combining RDC measurements and t2 analysis coupled with DFT calculations. A. Q factors of RDC measurements fitted for two possible stereoisomeric configurations of isoleucine. B. Q factors of RDC measurements fitted for four possible stereoisomeric configurations of xylose. C. Proton decoupled ^13^C spectrum of 65 mM xylose enantiomers in presence of 140 mg/ml (FK)_4_. D. T2 relaxation decay of xylose enantiomers. E. D-xylose interacts with (FK)_2_ model simulated with DFT. F. L-xylose interacts with (FK)_2_ model simulated with DFT. (FK)_2_ was represented with tube model, and xylose was represented with ball-stick model. Atoms are color coded respectively as red for O, blue for N, grey for C, and white for H; hydrogen bonding interactions are indicated by dashed lines.

After confirming the potential of (FK)_4_ on chiral differentiation to distinguish between enantiomers and identify absolute configurations, D-xylose was employed as another example. Xylose is a pentose sugar widely found in plants. It contains four hydroxyl functional groups, three of which exhibit intrinsic chirality, resulting in six possible configurations for xylose derivatives. Natural D-xylose exists in RSR configuration, while its enantiomer L-xylose has SRS chirality. A sample containing xylose enantiomers and the oligopeptide (FK)_4_ in D_2_O rapid self-assembled, and ready for NMR analysis within minutes. With (FK)_4_ functioning as the alignment media, RDC parameters were extracted from the [^1^H,^13^C]-J-scaled BIRD-HSQC (JSB-HSQC) spectra^28, 41^. A set of results ranging from 440 to 680 Hz were acquired (Table S2). Experimental RDC results of xylose were then used to fit DFT optimized structures of all four possible diastereomers using the StereoFitter program^28, 42^, with fitting quality assessed using quality factor n/x2 or Q value^28, 43-45^ (Figure 3B). The lowest Q value of 0.052 guided the assignment of RSR or its mirror-imaged SRS configuration among various diastereomers. To further differentiate between enantiomers, t2 measurements based on 1D proton-decoupled ^13^C spectra were applied to the same sample containing xylose enantiomers and (FK)_4_ used in the RDC experiment. Multiple chemical shifts demonstrated bifurcation, among which the diverged peak around 73 ppm corresponded to C3 (Figure 3C). Exponential decay fitting yielded distinct relaxation times for carbons in the two enantiomers (Figure 3D). T2 relaxation time of C3 is 1.05 s in D-xylose and is 0.64 s in the L-enantiomer, reflecting stronger interactions between L-xylose and (FK)_4_ around C3 atom compared to the D-enantiomer. DFT calculations monitored interactions between xylose enantiomers and a simplified model of (FK)_4_ oligopeptide. Both D- and L-xylose interacted with (FK)_4_ through hydrogen bonding at multiple hydroxyl group (OH) functionalized on the sugar ring and the ester oxygen (-O-) in the ring (Figure 3E and 3F). L-xylose exhibited a slightly stronger interaction with the oligopeptide compared to the D-xylose by 0.24 kcal/mol in electronic energy. One important aspect to note is that each enantiomer of xylose exists in inseparable α and β forms^46, 47^, distinguishable by two sets of split signals, such as peaks around 69.4 and 69.2 ppm for C4 (Figure 3C). Additionally, xylose can adopt several possible distributions in the pyranose ring, with the chair conformation being the most thermodynamically favorable^47^. Notably, a distinct environmental difference around C3 was observed in the simulation. In D-xylose, the hydroxyl group (OH) on C3 situated away from the oligopeptide without any interactions in the model (Figure 3E). On the other hand, this hydroxyl group (OH) hydrogen bounded directly to the amine group on the backbone of the residue in L-xylose (Figure 3F). The increased complexity of the chemical environment around C3 in L-xylose, compared to its D-enantiomer, was expected to result in a shorter t2 measurements. Consequently, the chemical shift at 73.33 ppm with a smaller t2 of 0.64 s was assigned to L-xylose, while the neighboring signal at 73.30 ppm was assigned to D-xylose.

### Chiral recognition and enantiopurity determination of various organic compounds

When oligopeptide (FK)_4_ acted as a chiral differentiating agent, bifurcation of chemical shifts corresponding to different enantiomers was observed in proton decoupled ^13^C NMR spectra, enabling assignment of absolute configurations. The peak differences for the same carbon between two enantiomers exceeded an average of 0.05 ppm, resulting in readily integrable individual peaks. This suggested the feasibility of chiral purity assessment in conjunction with simultaneous chiral recognition. Integration precision in conventional ^13^C NMR can be challenging due to the variable relaxation times of carbons with differing numbers of attached hydrogens^48^. However, similar chemical environments around identical carbon atoms in two enantiomers enable analyzable integration for chiral purity analysis. A sample containing 40 mM L-leucine and 25 mM of the corresponding D-enantiomer showed bifurcation of multiple signals in the presence of (FK)_4_ (Figure 4A). Each set of diverged resonances corresponding to the L- and D-enantiomers was sufficiently distinct for individual integration, resulting in a molar ratio close to 8:5, demonstrating the applicability of this method for quantifying enantiomeric purity. Furthermore, four additional samples with varying ratios of leucine enantiomers were analyzed (Figure 4B). Integration of peaks corresponding to each enantiomer, particularly focusing on branched resonance of C5 around 23 ppm, was achieved clearly. Ratios derived from ^13^C NMR integration were compared with the enantiomeric excess (e.e.) analysis via high-pressure liquid chromatography (HPLC), the most prevalent method for enantiomeric purity quantification. The high degree of linearity observed between these two analytical methods (Figure 4C) further supported the accuracy of chiral purity analysis and rapid chiral recognition using (FK)_4_ as a chiral differentiating agent.

**Figure 4.**
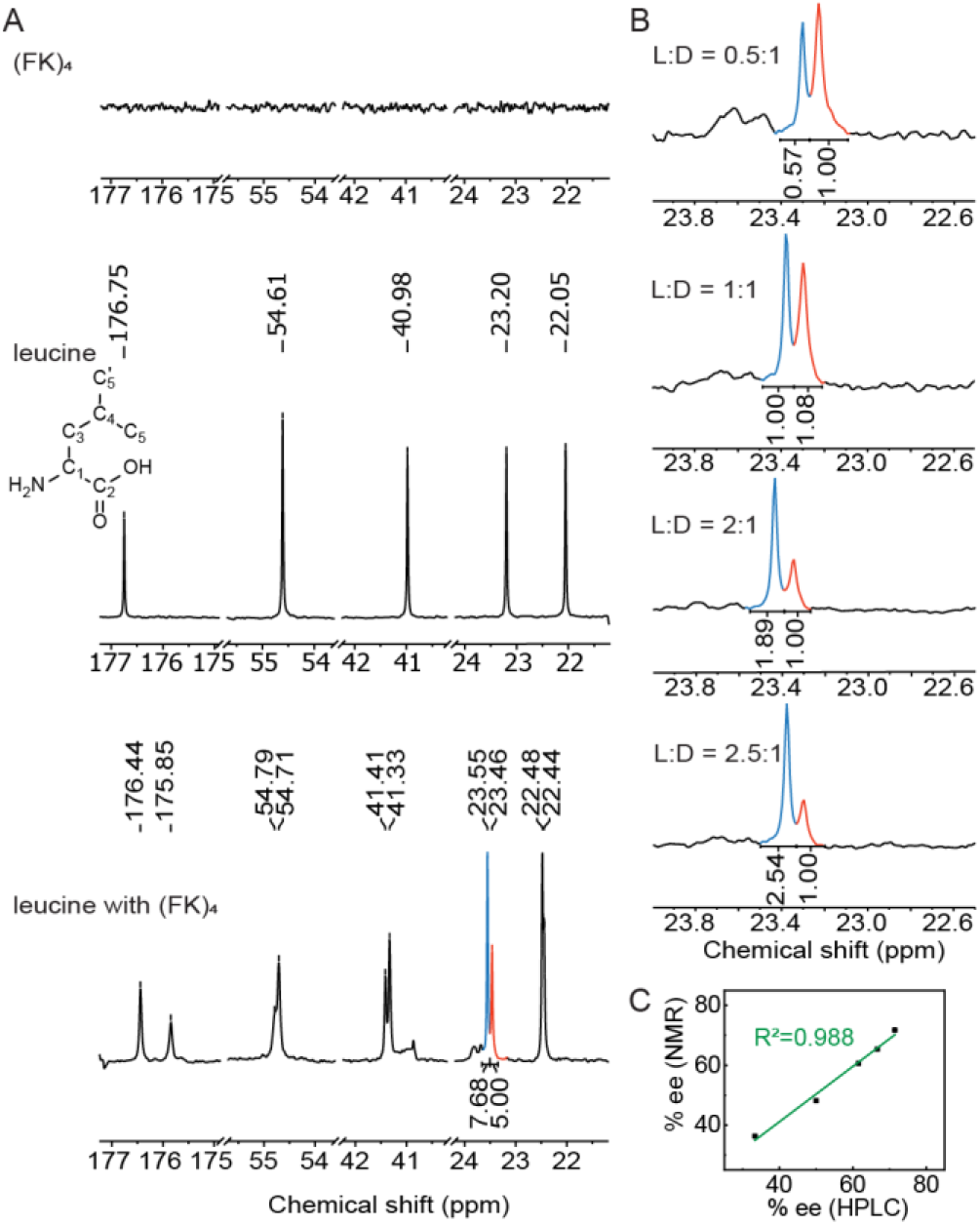
Chiral recognition and purity analysis of leucine enantiomers utilizing (FK)_4_ as the chiral differentiating agent in proton decoupled ^13^C NMR spectroscopy. A. Stacked spectrum of only (FK)_4_ (top), leucine enantiomers aqueous solution (middle), and leucine enantiomers aqueous solution mixed with the chiral differentiating agent (FK)_4_ (blue for L-leucine and red for D-leucine) (bottom). B. Integration of bifurcated peaks at around 22.90 ppm of the L- and D-enantiomers with presence of (FK)_4_. C. Linear correlation between the chiral purity analysis conducted via proton decoupled ^13^C NMR and HPLC.

To further demonstrate the practical utility of (FK)_4_ as a chiral differentiating agent, a diverse array of compounds spanning various types of organic molecules were subjected to chiral recognition *via* proton decoupled ^13^C NMR spectroscopy. Additional amino acids including proline, glutamic acid, glutamine, and phenylalanine exhibited distinct carbon resonance indicative of chiral recognition (Figure 5A to 5D). Notably, amino acids with positively charged side chains like arginine were not suitable for this method, possibly due to electrostatic repulsion between (FK)_4_ and the guanidinium group of arginine. Simple sugars like glucose, fructose, and galactose also demonstrated chiral recognition with (FK)_4_, evidenced by divergent carbon resonances (Figure 5E to 5G). Furthermore, biologically active organic acids and commercially available drugs including lactic acid, 2-methylvaleric acid, epinephrine, terbutaline, and chloroquine displayed successful chiral recognition with (FK)_4_ (Figure 5I to 5M). These examples underscored the broad applicability of (FK)_4_ as an effective chiral differentiating agent for a range of organic compounds across various molecular structures and functional groups in diverse chiral recognition analysis.

**Figure 5.**
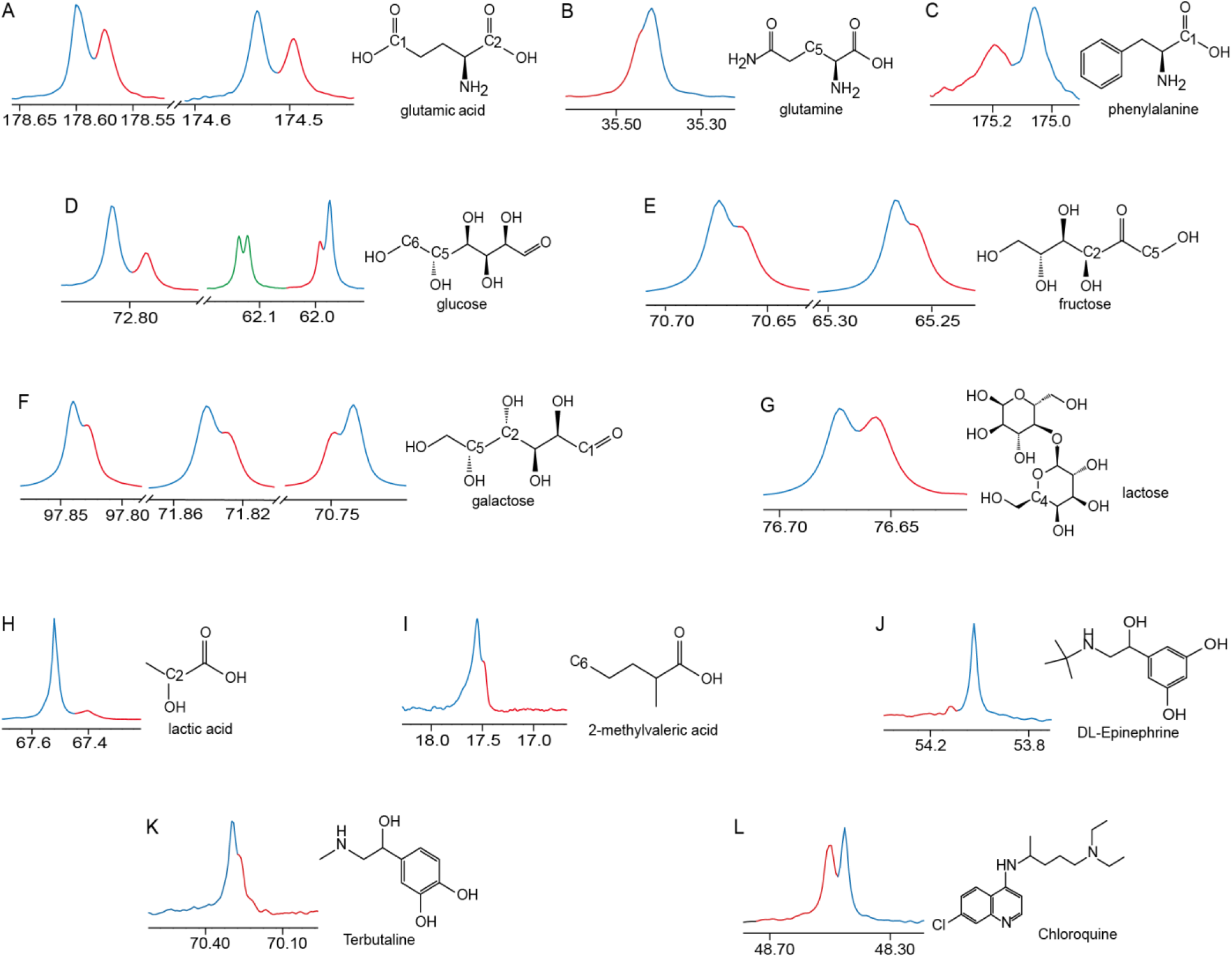
Chiral discrimination of diverse types of sample compounds with (FK)_4_ as a chiral differentiating agent using proton decoupled ^13^C NMR.

### Chiral recognition via ^1^H and ^19^F spectra

Broader types of NMR spectroscopy targeting different nuclei with higher sensitivity and reduced experimental time were tested with (FK)_4_ as chiral differentiating agent. ^1^H NMR is favored due to the high natural abundance and large gyromagnetic ratio of hydrogen nuclei^49^. Like ^13^C NMR, no recognizable peaks were observed in ^1^H NMR with bare (FK)_4_ (Figure 6A top). In the presence of (FK)_4_, difference between enantiomers of aspartic acid were detected in ^1^H NMR. The C3 proton peak at 3.00 ppm displayed as a complex multiple without (FK)_4_ (Figure 6A middle), but transformed into a set of symmetric splitting spectral lines upon addition of (FK)_4_ (Figure 6A bottom), indicating the presence of two enantiomers. The change of the resonance resembled the appearance of chemical shift non-equivalence induced by chiral interactions^18,22,50-51^. The resonance from aspartic acid at 3.00 ppm, split into symmetric two sets of three lines with varying intensities in the presence of (FK)_4_ (Figure 6A bottom), reflecting changes in the chemical shift due to chiral differentiation. The magnitude of splitting Δδ (change of chemical shift) correlated consistently with enantiomeric purity (Figure 6B). When L-aspartic acid was the predominant enantiomer, symmetric peaks emerge at 3.26 ppm and 2.81 ppm, with the splitting Δδ of 0.45 ppm (Figure 6B top). Conversely, when D-aspartic acid was the major enantiomer, symmetric peaks appear at 3.29 ppm and 2.78 ppm, with a Δδ of 0.51 ppm (Figure 6B bottom). Integration of these peaks in ^1^H NMR spectra provided quantitative information on enantiomer ratios, which aligned well with HPLC-based enantiomeric excess analysis (Figure 6D). ^19^F NMR is also extensively used in organic synthesis due to its high sensitivity and robust chemical shift dispersion^52, 53^. 4-fluorophenylethanamine was selected as the analyte to evaluate the effectiveness of (FK)_4_ as a chiral differentiating agent in ^19^F NMR. An aqueous sample, comprising both L- and D-enantiomer, exhibited one single fluorene signal at −113.34 ppm in the proton decoupled ^19^F spectra (Figure 6C top). Upon addition of (FK)_4_, this singlet peak diverged into two distinct peaks at −113.45 ppm and −113.58 ppm with a chemical shift difference of 0.13 ppm (Figure 6C bottom). Integration of each peak resulted in an 8:5 ratio of the L-to D-enantiomer, consistent with HPLC-based enantiomeric ratio determination. These results highlighted the broad applicability of (FK)_4_ as chiral differentiating agent in diverse contexts for pivotal chiral analysis.

**Figure 6.**
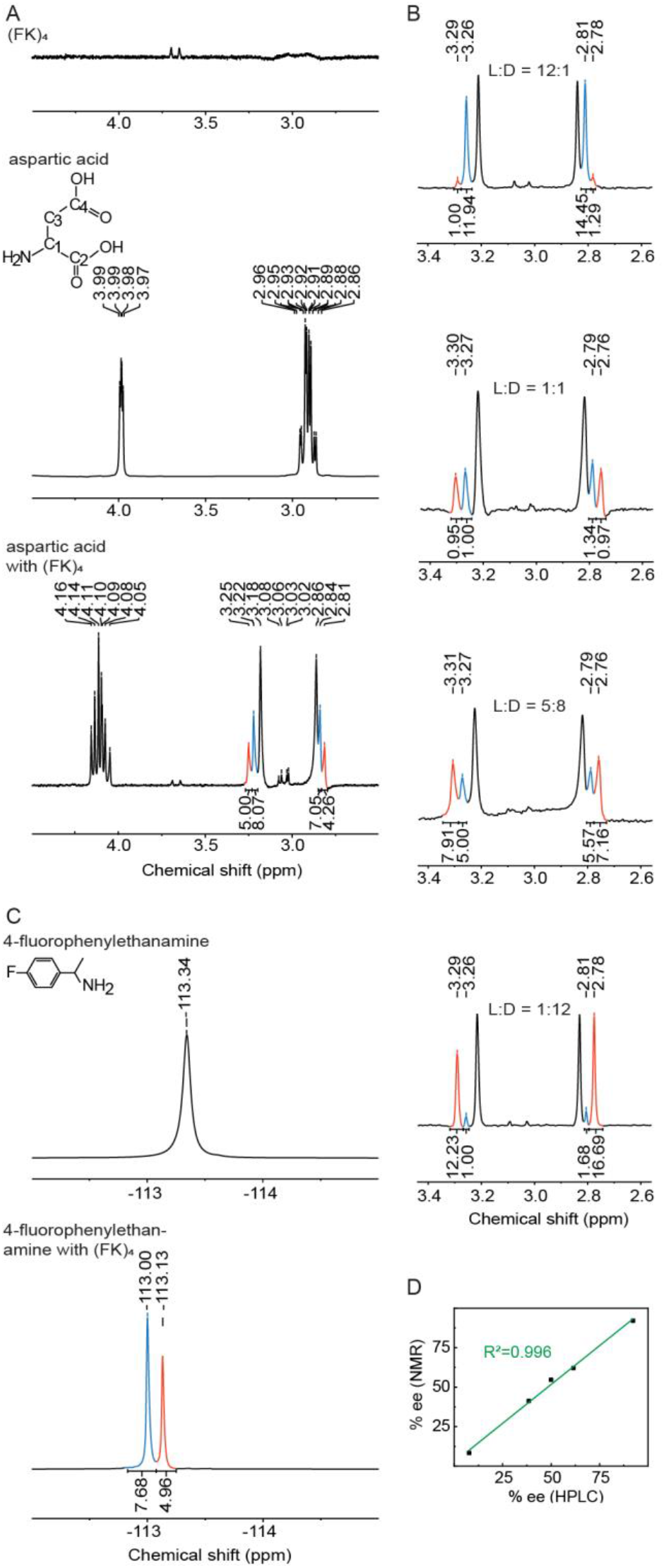
Chiral recognition and purity analysis of an aspartic acid enantiomeric sample and a 4-fluorophenylethanamine enantiomeric sample utilizing (FK)_4_ as the chiral differentiating agent in ^1^H and ^19^F NMR spectroscopy respectively. A. Stacked spectrum of only (FK)_4_ (top), aspartic acid enantiomers aqueous solution (middle), and aspartic acid enantiomers aqueous solution mixed with the chiral differentiating agent (FK)_4_ (blue for L-aspartic acid and red for D-aspartic acid) (bottom). B. Integration of the symmetric bifurcated resonance at around 3 ppm of the Land D-enan- tiomers with presence of (FK)_4_. C. Stacked spectrum of 4-fluorophenylethanamine aqueous solution (top) and the aqueous solution with presence of (FK)_4_ (blue for L-enantiomer and red for D-enantiomer) (bottom). D. Linear correlation between the chiral purity analysis of aspartic acid enantiomers conducted via ^1^H NMR and HPLC.

## Conclusion

In this study, we investigated the versatile application of oligopeptide (FK)_4_, which exhibited dual properties as both an alignment media and a chiral differentiating agent. Function as an alignment media, (FK)_4_ was effective for anisotropic NMR measurements such as RDC, facilitating structural elucidation of stereoisomers. Function as a chiral differentiating agent, (FK)_4_ interacted with enantiomers uniquely, resulting in bifurcation of chemical shift signals, particularly evident for proton decoupled carbons. By demonstrating this application with isoleucine and xylose, the culmination of the dual functionality of (FK)_4_ enabled the identification of stereoisomeric structure *via* RDC parameters and assignment of enantiomeric structure *via* t2 measurements coupled with theoretical simulations. This utilization of oligopeptide (FK)_4_ advanced the elucidation of absolute configurations of organic molecules. Our findings underscored the potential of (FK)_4_ as a valuable tool in both stereoisomeric elucidation and enantiomer discrimination.

## Supporting information

Experimental details, cryo-electron microscopy results, RDC data, computational details, and additional NMR results.

## Notes

The authors declare no competing financial interest.

## ACKNOWLEDGMENT

The authors thank NMR facility faculties at the Innovation Academy of Precision Measurement Science and Technology for their assistance in maintenance and utilization of NMR instruments. The authors thank the funding sources assisted in this work.

## ABBREVIATIONS

NMR: nuclear magnetic resonance
CDA: chiral derivatization agent
CSA: chiral solvating agent
TEM: transmission electron microscopy; e.e., enantiomeric excess

