## Supplementary material for "Oligopeptide (FK)_4_ facilitates the determination of the absolute stereochemistry of chiral organic compounds *via* NMR Spectroscopy": Experimental details, cryo-electron microscopy results, RDC data, computational details, and additional NMR results.

#### 1. Materials and Methods

##### 1.1 Self-assembly sample preparation

L- and D-amino acids and other analytes are weighed in proportion and dissolved in $D_2O$. Adjust the solution to pH=3 with hydrochloric acid. $(FK)_4$ is dissolved in the mixture, vortexed, and centrifuged at 4000g for 1 minute to remove bubbles. This process is repeated until $(FK)_4$ is completely dissolved. The self-assembly process takes less than 10 minutes. The liquid can be directly transferred to a nuclear magnetic tube and sampled in a magnetic field.

##### 1.2 TEM

For negative staining, 300-mesh holey carbon grids were glow-discharged at 15mA for 25s before use. An aliquot of 4 μl of the sample was loaded onto grids for 30s, followed by blotting and staining with 2% uranyl acetate. Grids were then transferred onto Talos L120C transmission electron microscope at 120kV equipped with 4k × 4k Ceta camera and images were recorded at 92000× with a defocus range of -2.0 to -3.0 μm.

##### 1.3 Cryo-TEM

For cryo-EM datasets collection, 3 μl of the samples at a concentration of 10mg/ml was applied to glow-discharged Cu grids (Quantifoil, R1.2/1.3, 300-mesh). The grids were blotted for 3s and then plunge-frozen into liquid ethane using Vitrobot Mark IV (Themo Fischer). Micrographs were recorded using 4k×4k CetaD camera with EPU software on Glacios TEM at 200kV. Images were obtained at 120000× with a pixle size of 1.2 Å and a defocus of -3 μm. The exposure time was 1.89s, resulting a total dose of 34e/ $Å^2$.

##### 1.4 NMR Experiments

If not specified, all NMR experiments for $(FK)_4$ and $(FK)_4$-additives complexes were performed in $D_2O$ and the concerntration of $(FK)_4$ is 140mg/ml. If not specified, all spectra were collected at 298 K by Bruker Avance 600 NMR spectrometers equipped with cryogenically cooled probes. The NMR data were processed in NMRPipe and analyzed in MestReNova[1].

##### 1.5 RDC measurements

F2-coupled [$^{1}$H,$^{13}$C]-CLIP-HSQC spectra were acquired with the number of scans 8, the relaxation delay of 2 s, data points of 8192 ($t_1$) x 256 ($t_2$), and spectral widths of 14 ppm for $^{1}$H and 180 ppm for $^{13}$C. F1-coupled [$^{1}$H,$^{13}$C]-JSB-HSQC spectra were acquired with the number of scans 8, the relaxation delay of 2 s, data points of 2048 ($t_1$) x 1024 ($t_2$), J-scaling factor of 4, and spectral widths of 14 ppm for $^{1}$H and 180 ppm for $^{13}$C. Errors of proton-carbon coupling were estimated following published methods[2, 3, 4].

Configuration for analytes were drawn in ChemDraw. The initial energy-minimization of configurations of analytes were done by ChemDraw 3D. Diastereoisomers were generated by MestReNova software based on the

2D structure of the analytes. Conformers of each possible diastereoisomers were generated in Macromodel4 using the torsional sampling (MCMM) method with the OPLS_2005 force-field and the energy threshold of 5 kcal/mol. DFT geometry optimization for each conformer was performed in Gaussian 095 using the B3LYP functional and the 6-31G** basis. Fitting of RDC data to each possible diastereoisomers was done by the StereoFitter[5] program.

##### 1.6 The T2 spectrum

The NMR measures the relaxation signals of carbon atoms, and the Carr-Purcell-Meiboom-Gill (CPMG) pulse sequences combined with inverse gated-decoupling is used to represent the signals. Through the attenuation process of echo signals, the transverse relaxation time ($T_2$ time) can be calculated, and the echo amplitude decays exponentially at the rate of $1/T_2$. The $T_2$ spectrum is represented by 10 logarithmic evenly distributed data points and this process can be fitted with the exponential formula:

$$f(t) = A_1 e^{-t/T_2} \quad (1)$$

The $T_2$ spectrum can be obtained by exponential inversion fitting of the carbon atom spin echoes using formula (1). In order to improve the signal-to-noise ratio (SNR) during the inversion of the $T_2$ spectra, the $T_2$ spectrum can be accumulated 32 times in the depth domain to improve the accuracy of $T_2$ spectral inversion.

##### 1.7 DFT

DFT calculations simulating interactions between analytes and $(FK)_2$ model. All calculations were performed with Gaussian16 using density functional theory with B3LYP hybrid functional with D3 dispersion corrections was used for all calculations. All atoms (H, C, N, and O) were treated with the 6-31++G(d,p) basis set. The solvent ($H_2O$) was modeled by the solvation method based on density (SMD). All geometry optimizations were carried out in solution.

#### 2. Cryo-Electron Microscopy

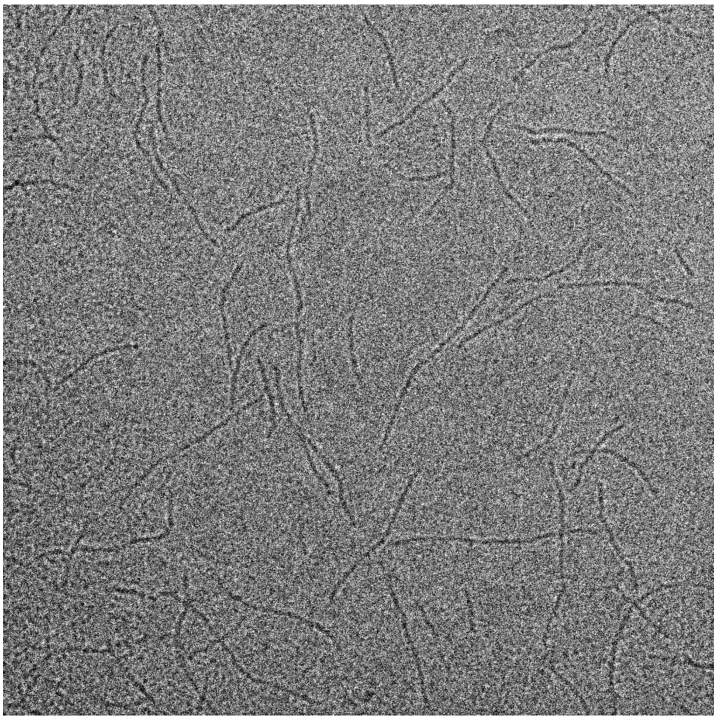

Figure S1. Cryo-electron microscopy image of self-assembled oligopeptide $(FK)_4$.

#### 3. RDC data

##### 3.1 RDC data of isoleucine

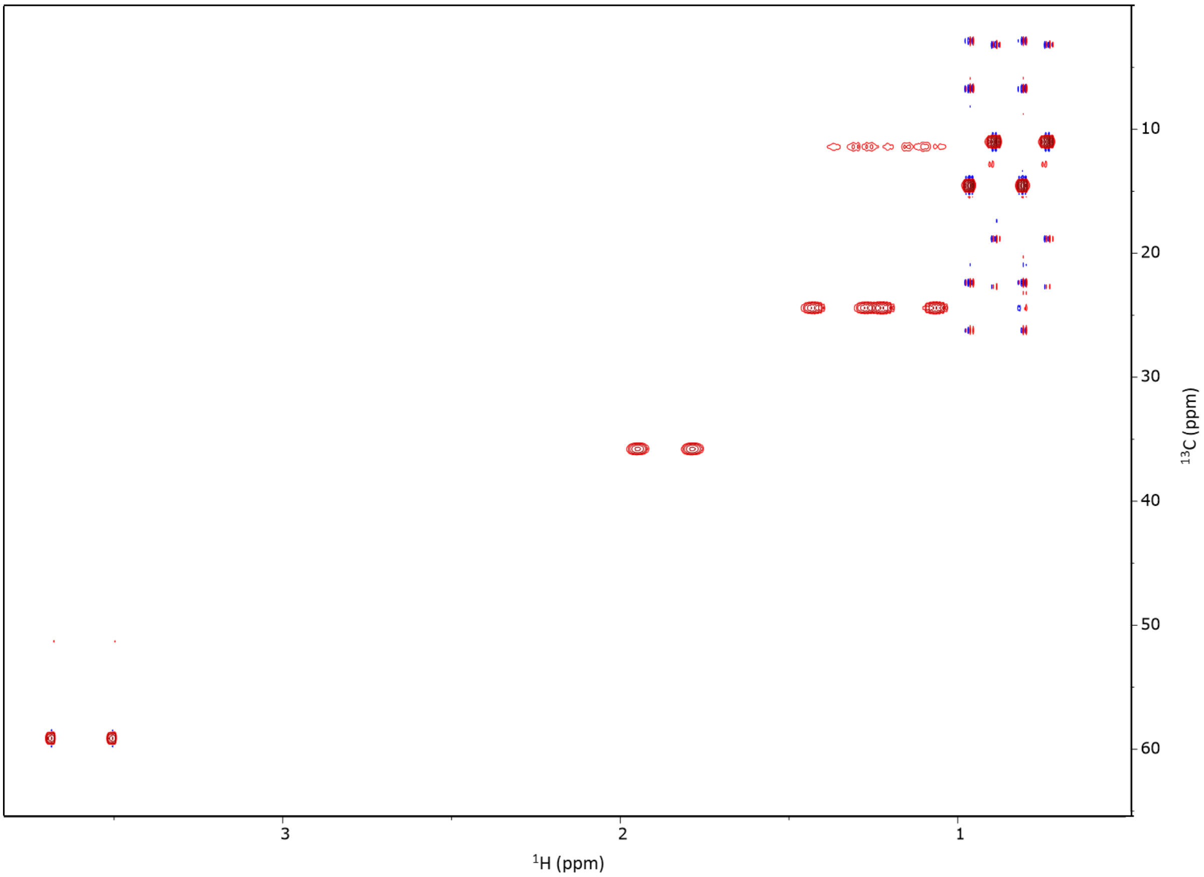

Figure S2. The overlaid [$^{1}$H, $^{13}$C]-JSB-HSQC spectra of isoleucine in the isotropic $D_2O$ (pink and purple) and anisotropic $(FK)_4/D_2O$ (cyan and orange).

| Atom number | $^{1}J_{CH}$ | $^{1}T_{CH}$ | $^{1}D_{CH}$ | $^{1}D_{CH}$ calculated |
|---|---|---|---|---|
| C3/H3 | 144.8 | 377.95 | 233.15 | 233.147 |
| C4/H4 | 132.1 | 386.69 | 254.59 | 254.615 |
| C5/H5$_{a,b}$ | 25.8 | 55.02 | 14.61 | 14.116 |
| C6/H6a | 124.9 | 134 | 9.1 | 9.191 |
| C6/H6b | 125 | 122.2 | -2.8 | -2.527 |
| C6/H6c | 125.8 | 129.5 | 3.7 | 4.013 |
| C7/H7 Me | 126.8 | 126.7 | -0.1 | -0.057 |

Table S1. Proton-carbon J-coupling ($^{1}J_{CH}$), proton-carbon total coupling ($^{1}T_{CH}$), experimental RDC values ($^{1}D_{CH}$), and calculated RDC values ($^{1}D_{CH}$ calculated) of isoleucine with 140 mg/ml $(FK)_4$.

#### 3.2 RDC data of xylose

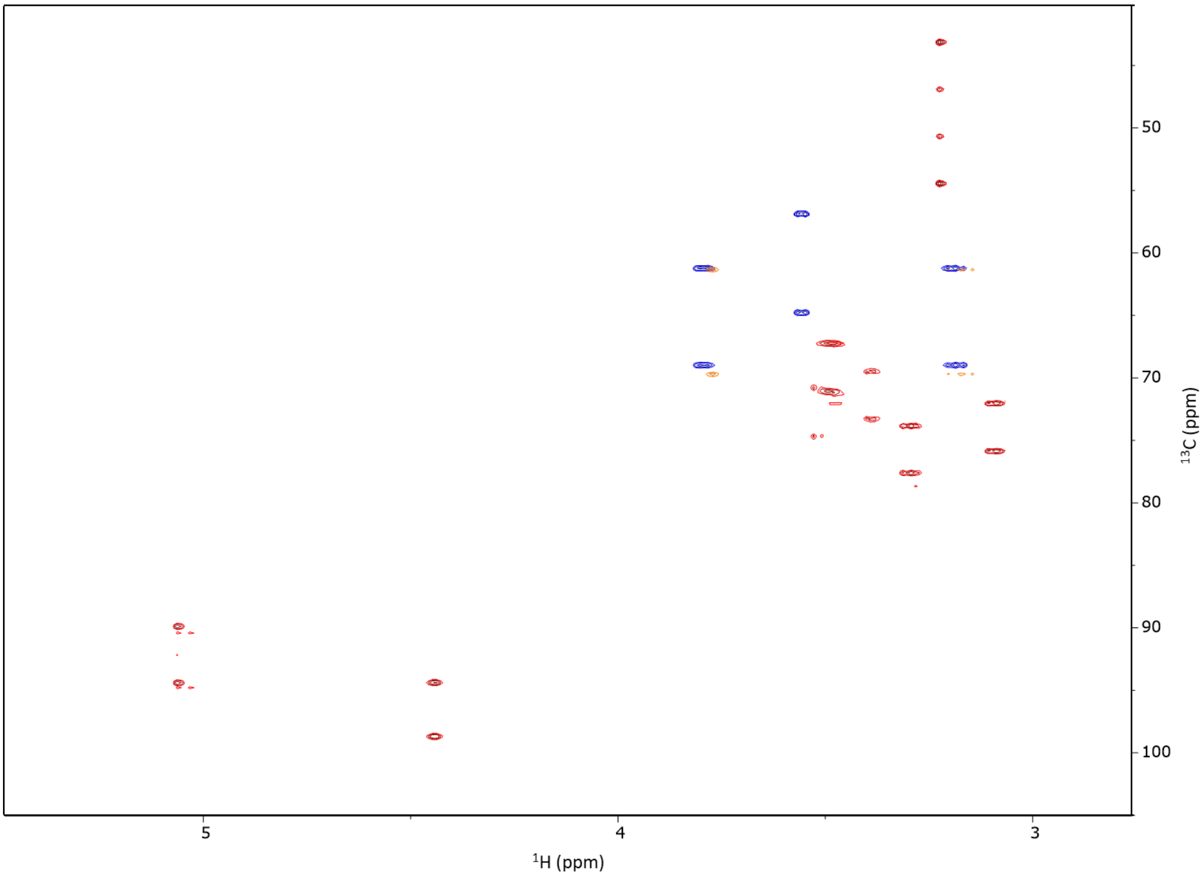

Figure S3. The overlaid [$^1$H, $^{13}$C]-JSB-HSQC spectra of xylose in the isotropic $D_2O$ (pink and purple) and anisotropic $(FK)_4/D_2O$ (cyan and orange).

| Atom number | $^1J_{CH}$ | $^1T_{CH}$ | $^1D_{CH}$ | $^1D_{CH}$ calculated |
|---|---|---|---|---|
| C1/H1$_{a,b}$ | 442.96 | 499.61 | 56.657 | 55.697 |
| C2/H2 | 576.87 | 721.09 | 144.218 | 143.962 |
| C3/H3 | 556.27 | 762.3 | 206.026 | 214.623 |
| C4/H4 | 576.87 | 762.3 | 185.423 | 181.75 |
| C5/H5 | 679.89 | 659.28 | -20.603 | -7.038 |

Table S2. Proton-carbon J-coupling ($^1J_{CH}$), proton-carbon total coupling ($^1T_{CH}$), experimental RDC values ($^1D_{CH}$), and calculated RDC values ($^1D_{CH}$ calculated) of xylose with 140 mg/ml $(FK)_4$.

##### 3.3 Input files of isoleucine for StereoFitter

```
rdc_data {
# C5 H5a,b
 (5,14) (5,15) 14.610
# C4 H4
 4 13 254.590
# C3 H3
 3 12 233.150
# C6 H6a
 6 16 9.1
# C6 H6b
 6 17 -2.8
# C6 H6c
 6 18 3.7
# C7 H7 Me
 (7,19) (7,20) (7,21) -0.1
}

rdc_standard_error {
 1.2
}

superimpose {
 true
}

superimpose_atoms {
 auto
}
```

##### 3.4 Input files of xylose for StereoFitter

```
rdc_data {
# C1 H1a,b
 (1,11) (1,12)  56.657
# C2 H2
 2 13 144.218
# C3 H3
 3 14 206.026
# C4 H4
 4 15 185.423
# C5 H5
 5 16 -20.603
}

rdc_standard_error {
 1.2
}

superimpose {
 true
}

superimpose_atoms {
 auto
}
```

### 4. DFT model of isoleucine with $(FK)_4$

#### 4.1 DFT model of isoleucine with oligopeptide $(FK)_2$ model

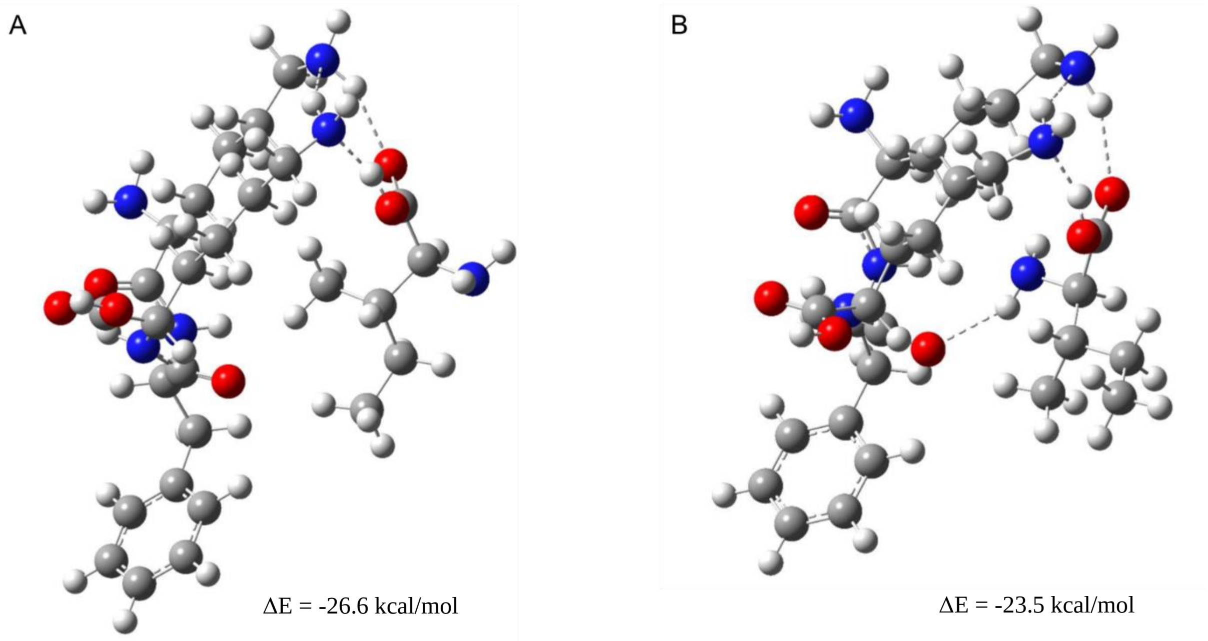

Figure S4. Comparison of interactions between isoleucine enantiomers with $(FK)_4$ investigated with DFT calculation. A. L-isoleucine interacts with $(FK)_4$ model simulated with DFT. B. D-isoleucine interacts with $(FK)_4$ model simulated with DFT. Atoms are color coded respectively as red for O, blue for N, grey for C, and white for H; hydrogen bonding interactions are indicated by dashed lines.

#### 4.2 Coordinates of all interacting structures between $(FK)_2$ model and analytes

| Atomic Number | Coordinates (Angstroms) | | |
|---|---|---|---|
| | X | Y | Z |
| 6 | -16.4497 | 10.198 | 9.529362 |
| 6 | -15.2115 | 9.53163 | 10.11315 |
| 6 | -17.7052 | 9.487601 | 10.08396 |
| 8 | -14.757 | 8.4784 | 9.630851 |
| 6 | -18.9516 | 10.3734 | 10.00956 |
| 6 | -20.2175 | 9.67819 | 10.51895 |
| 6 | -21.3779 | 10.64837 | 10.71647 |
| 7 | -14.6819 | 10.14039 | 11.18955 |
| 6 | -13.5521 | 9.607694 | 11.94895 |
| 6 | -14.0898 | 8.825604 | 13.1653 |
| 6 | -12.6019 | 10.74371 | 12.34982 |
| 8 | -13.9426 | 9.217258 | 14.32662 |
| 6 | -11.2997 | 10.26256 | 12.95236 |
| 6 | -10.9367 | 10.60603 | 14.25875 |

| | | | |
|---|---|---|---|
| 6 | -10.4204 | 9.470846 | 12.20177 |
| 6 | -9.72857 | 10.1703 | 14.80479 |
| 6 | -9.21408 | 9.029699 | 12.74426 |
| 6 | -8.86361 | 9.377692 | 14.05004 |
| 7 | -14.7562 | 7.699839 | 12.83597 |
| 6 | -15.5771 | 6.937651 | 13.76468 |
| 6 | -15.4743 | 5.463321 | 13.42536 |
| 6 | -17.0597 | 7.380639 | 13.7104 |
| 8 | -15.1115 | 5.01762 | 12.35577 |
| 6 | -17.2987 | 8.721994 | 14.41292 |
| 6 | -18.6204 | 9.361327 | 13.98666 |
| 6 | -18.9983 | 10.5416 | 14.86966 |
| 1 | -17.5349 | 9.211956 | 11.12879 |
| 1 | -17.8511 | 8.553467 | 9.529333 |
| 1 | -19.1116 | 10.71692 | 8.980246 |
| 1 | -18.7659 | 11.26898 | 10.61384 |
| 1 | -20.004 | 9.193341 | 11.47772 |
| 1 | -20.5152 | 8.881579 | 9.827349 |
| 1 | -21.6975 | 11.04652 | 9.748797 |
| 1 | -21.0239 | 11.50496 | 11.30785 |
| 1 | -16.4712 | 11.23436 | 9.867393 |
| 1 | -13.1153 | 11.4148 | 13.04188 |
| 1 | -12.3906 | 11.31411 | 11.44132 |
| 1 | -11.6078 | 11.21583 | 14.85399 |
| 1 | -10.6807 | 9.199902 | 11.18349 |
| 1 | -9.46553 | 10.44779 | 15.81975 |
| 1 | -8.54686 | 8.417738 | 12.14707 |
| 1 | -7.92565 | 9.035876 | 14.47323 |
| 1 | -13.032 | 8.903025 | 11.29891 |
| 1 | -15.2079 | 10.91103 | 11.61091 |
| 1 | -17.6849 | 6.611131 | 14.17043 |
| 1 | -17.3463 | 7.442703 | 12.65576 |
| 1 | -16.4955 | 9.42467 | 14.18072 |
| 1 | -17.2759 | 8.565091 | 15.49712 |
| 1 | -19.4331 | 8.62706 | 13.99704 |
| 1 | -18.5147 | 9.709921 | 12.95774 |
| 1 | -18.1334 | 11.16598 | 15.09672 |
| 1 | -19.4507 | 10.21804 | 15.80633 |
| 1 | -15.186 | 7.080147 | 14.77258 |
| 1 | -14.8791 | 7.490158 | 11.85124 |
| 7 | -22.5342 | 9.97842 | 11.34289 |
| 1 | -22.2708 | 9.692979 | 12.28428 |
| 1 | -23.2875 | 10.65113 | 11.45966 |
| 7 | -19.9784 | 11.43149 | 14.15874 |
| 1 | -20.6825 | 10.8892 | 13.65263 |
| 1 | -20.4637 | 12.046 | 14.81415 |
| 8 | -15.8778 | 4.692806 | 14.4484 |

| | | | |
|---|---|---|---|
| 1 | -15.8562 | 3.759593 | 14.17541 |
| 7 | -16.3385 | 10.21032 | 8.063062 |
| 1 | -16.3157 | 9.246377 | 7.736756 |
| 1 | -17.1886 | 10.61695 | 7.681198 |
| 8 | -16.6612 | 11.77596 | 12.46107 |
| 6 | -17.0963 | 12.90408 | 12.8111 |
| 6 | -16.0489 | 13.98628 | 13.12678 |
| 1 | -15.1789 | 13.44385 | 13.51745 |
| 8 | -18.3292 | 13.19408 | 12.90758 |
| 1 | -19.445 | 12.04177 | 13.48499 |
| 6 | -15.5795 | 14.72844 | 11.84557 |
| 1 | -14.9798 | 15.57068 | 12.21488 |
| 7 | -16.5622 | 14.9615 | 14.10558 |
| 1 | -16.928 | 14.46382 | 14.91271 |
| 1 | -15.778 | 15.51526 | 14.44095 |
| 6 | -14.6686 | 13.8615 | 10.97058 |
| 1 | -15.2239 | 13.03495 | 10.52127 |
| 1 | -14.2471 | 14.45898 | 10.15663 |
| 1 | -13.8404 | 13.43811 | 11.54519 |
| 6 | -16.7377 | 15.29746 | 11.01797 |
| 1 | -16.3521 | 15.92684 | 10.21034 |
| 1 | -17.3229 | 14.4928 | 10.56079 |
| 1 | -17.4122 | 15.90328 | 11.62647 |

Table S3. Interacting structure between L-valine and $(FK)_2$ model

| Atomic Number | Coordinates (Angstroms) | | |
|---|---|---|---|
| | X | Y | Z |
| 6 | -16.102233 | 10.684066 | 9.904893 |
| 6 | -14.994703 | 9.838128 | 10.518843 |
| 6 | -17.438504 | 10.215131 | 10.518478 |
| 8 | -14.741353 | 8.70054 | 10.069836 |
| 6 | -18.636837 | 11.07668 | 10.112683 |
| 6 | -19.93182 | 10.609733 | 10.783514 |
| 6 | -21.094449 | 11.575204 | 10.591654 |
| 7 | -14.372425 | 10.360388 | 11.583632 |
| 6 | -13.319599 | 9.64651 | 12.310708 |
| 6 | -13.957271 | 8.718836 | 13.368568 |
| 6 | -12.330298 | 10.643171 | 12.926261 |
| 8 | -13.89589 | 8.949347 | 14.580764 |
| 6 | -11.086519 | 9.983534 | 13.482259 |
| 6 | -10.781949 | 10.046902 | 14.845877 |
| 6 | -10.205616 | 9.301771 | 12.632208 |
| 6 | -9.628623 | 9.444441 | 15.350629 |
| 6 | -9.054091 | 8.695353 | 13.132274 |
| 6 | -8.76151 | 8.763991 | 14.495692 |
| 7 | -14.593121 | 7.655807 | 12.837424 |

| | | | |
|---|---|---|---|
| 6 | -15.432664 | 6.739679 | 13.59232 |
| 6 | -15.285737 | 5.344958 | 13.014519 |
| 6 | -16.923984 | 7.155628 | 13.555393 |
| 8 | -14.904827 | 5.092334 | 11.889523 |
| 6 | -17.221551 | 8.335522 | 14.48327 |
| 6 | -18.603834 | 8.938684 | 14.226312 |
| 6 | -18.992297 | 9.939219 | 15.312239 |
| 1 | -17.341271 | 10.239195 | 11.608936 |
| 1 | -17.607026 | 9.169563 | 10.233766 |
| 1 | -18.767083 | 11.063952 | 9.024516 |
| 1 | -18.434069 | 12.115626 | 10.395766 |
| 1 | -19.755956 | 10.498177 | 11.857682 |
| 1 | -20.209802 | 9.618175 | 10.407423 |
| 1 | -21.354179 | 11.643343 | 9.530845 |
| 1 | -20.774087 | 12.57852 | 10.906528 |
| 1 | -15.945885 | 11.728506 | 10.182516 |
| 1 | -12.831503 | 11.219914 | 13.705937 |
| 1 | -12.044534 | 11.341344 | 12.134288 |
| 1 | -11.455333 | 10.568601 | 15.517194 |
| 1 | -10.421957 | 9.248102 | 11.569903 |
| 1 | -9.410011 | 9.504948 | 16.411226 |
| 1 | -8.384183 | 8.172879 | 12.45809 |
| 1 | -7.866173 | 8.293492 | 14.886474 |
| 1 | -12.79727 | 9.01534 | 11.589203 |
| 1 | -14.645118 | 11.307516 | 11.917669 |
| 1 | -17.544076 | 6.302063 | 13.84262 |
| 1 | -17.175218 | 7.40681 | 12.51936 |
| 1 | -16.476577 | 9.12382 | 14.347748 |
| 1 | -17.137928 | 7.998847 | 15.522929 |
| 1 | -19.366124 | 8.155737 | 14.157973 |
| 1 | -18.580636 | 9.44272 | 13.257755 |
| 1 | -18.119212 | 10.47331 | 15.685028 |
| 1 | -19.490063 | 9.453859 | 16.15087 |
| 1 | -15.084494 | 6.715126 | 14.625405 |
| 1 | -14.660503 | 7.612582 | 11.823109 |
| 7 | -22.286858 | 11.112896 | 11.329937 |
| 1 | -22.064227 | 11.096024 | 12.323601 |
| 1 | -23.032771 | 11.795699 | 11.2252 |
| 7 | -19.909348 | 10.991834 | 14.765007 |
| 1 | -20.685668 | 10.579242 | 14.243447 |
| 1 | -20.301371 | 11.564638 | 15.513547 |
| 8 | -15.672514 | 4.405028 | 13.892387 |
| 1 | -15.624474 | 3.530031 | 13.470358 |
| 7 | -16.011128 | 10.589196 | 8.440442 |
| 1 | -16.163017 | 9.619355 | 8.171465 |
| 1 | -16.775722 | 11.119166 | 8.032231 |
| 8 | -16.650577 | 12.282787 | 14.544466 |

| | | | |
|---|---|---|---|
| 6 | -17.176672 | 12.721226 | 13.490522 |
| 6 | -16.319005 | 13.59842 | 12.563131 |
| 8 | -18.384683 | 12.529145 | 13.143954 |
| 1 | -19.349417 | 11.623438 | 14.112425 |
| 7 | -14.953178 | 13.045408 | 12.514435 |
| 1 | -14.354546 | 13.652002 | 11.961358 |
| 1 | -14.570251 | 13.037221 | 13.457841 |
| 1 | -16.744835 | 13.536932 | 11.557781 |
| 6 | -16.374594 | 15.08328 | 13.027008 |
| 1 | -15.874672 | 15.124133 | 14.003485 |
| 6 | -15.612638 | 15.988455 | 12.049707 |
| 1 | -15.641168 | 17.026937 | 12.392009 |
| 1 | -14.560261 | 15.709483 | 11.950303 |
| 1 | -16.070139 | 15.949749 | 11.054813 |
| 6 | -17.810998 | 15.59528 | 13.194363 |
| 1 | -18.369701 | 15.498479 | 12.257353 |
| 1 | -18.356421 | 15.050834 | 13.968137 |
| 1 | -17.803222 | 16.653307 | 13.47208 |

Table S4. Interacting structure between D-valine and $(FK)_2$ model

| Atomic Number | Coordinates (Angstroms) | | |
|---|---|---|---|
| | X | Y | Z |
| 6 | -15.274905 | 9.71794 | 8.82769 |
| 6 | -14.190029 | 9.171181 | 9.744172 |
| 6 | -16.618557 | 9.555288 | 9.570403 |
| 8 | -13.996902 | 7.951082 | 9.850764 |
| 6 | -17.790101 | 10.257576 | 8.88104 |
| 6 | -19.070047 | 10.18534 | 9.717754 |
| 6 | -20.253464 | 10.889948 | 9.065076 |
| 7 | -13.498027 | 10.085701 | 10.460268 |
| 6 | -12.640761 | 9.724633 | 11.592566 |
| 6 | -13.538297 | 9.700056 | 12.842478 |
| 6 | -11.472268 | 10.705581 | 11.733642 |
| 8 | -13.708847 | 10.701933 | 13.543999 |
| 6 | -10.521965 | 10.302676 | 12.839659 |
| 6 | -10.435259 | 11.04169 | 14.023967 |
| 6 | -9.721353 | 9.161211 | 12.701314 |
| 6 | -9.569324 | 10.652103 | 15.047018 |
| 6 | -8.857093 | 8.767103 | 13.721689 |
| 6 | -8.778924 | 9.512223 | 14.900031 |
| 7 | -14.167454 | 8.527888 | 13.065122 |
| 6 | -15.290579 | 8.436175 | 13.99239 |
| 6 | -15.526345 | 6.977749 | 14.322869 |
| 6 | -16.560101 | 9.068145 | 13.373519 |
| 8 | -15.26054 | 6.054625 | 13.578583 |
| 6 | -17.719708 | 9.303001 | 14.342231 |

| | | | |
|---|---|---|---|
| 6 | -18.798355 | 10.155579 | 13.669874 |
| 6 | -19.932231 | 10.523186 | 14.61326 |
| 1 | -16.510653 | 9.958327 | 10.580585 |
| 1 | -16.827217 | 8.484345 | 9.676545 |
| 1 | -17.970567 | 9.815465 | 7.894086 |
| 1 | -17.528251 | 11.309022 | 8.712761 |
| 1 | -18.883885 | 10.645594 | 10.693623 |
| 1 | -19.330689 | 9.137717 | 9.910631 |
| 1 | -20.53716 | 10.364505 | 8.148223 |
| 1 | -19.944678 | 11.902522 | 8.769008 |
| 1 | -15.098384 | 10.783491 | 8.661022 |
| 1 | -11.862661 | 11.708704 | 11.916107 |
| 1 | -10.944209 | 10.721986 | 10.776557 |
| 1 | -11.052147 | 11.925258 | 14.145703 |
| 1 | -9.77591 | 8.579449 | 11.786536 |
| 1 | -9.514497 | 11.237982 | 15.958042 |
| 1 | -8.243529 | 7.881595 | 13.596822 |
| 1 | -8.107007 | 9.20745 | 15.694615 |
| 1 | -12.257356 | 8.72315 | 11.400798 |
| 1 | -13.797262 | 11.052368 | 10.407214 |
| 1 | -16.880677 | 8.454309 | 12.526526 |
| 1 | -16.244568 | 10.031484 | 12.96737 |
| 1 | -17.348133 | 9.82268 | 15.23281 |
| 1 | -18.143676 | 8.352748 | 14.679982 |
| 1 | -19.204477 | 9.629187 | 12.800016 |
| 1 | -18.342049 | 11.077975 | 13.300837 |
| 1 | -19.556307 | 11.006519 | 15.515798 |
| 1 | -20.525154 | 9.655013 | 14.901901 |
| 1 | -15.031257 | 8.966647 | 14.908557 |
| 1 | -14.088012 | 7.801867 | 12.361837 |
| 7 | -21.42114 | 10.907389 | 9.968286 |
| 1 | -21.193833 | 11.503679 | 10.762018 |
| 1 | -22.199668 | 11.362174 | 9.498387 |
| 7 | -20.843368 | 11.510917 | 13.947144 |
| 1 | -21.229494 | 11.124121 | 13.08325 |
| 1 | -21.624758 | 11.765379 | 14.552826 |
| 8 | -16.105052 | 6.817688 | 15.520729 |
| 1 | -16.293939 | 5.873491 | 15.662605 |
| 7 | -15.189183 | 9.036902 | 7.52796 |
| 1 | -15.360338 | 8.04429 | 7.676192 |
| 1 | -15.948924 | 9.368083 | 6.939858 |
| 8 | -19.59552 | 13.437523 | 11.292409 |
| 6 | -18.922263 | 13.859905 | 12.267676 |
| 6 | -17.680637 | 14.735603 | 11.993705 |
| 1 | -17.728628 | 15.039982 | 10.944394 |
| 8 | -19.19508 | 13.612723 | 13.488599 |
| 1 | -20.290551 | 12.388055 | 13.704193 |

| | | | |
|---|---|---|---|
| 7 | -17.672841 | 15.957967 | 12.818938 |
| 1 | -18.547467 | 16.455795 | 12.671388 |
| 1 | -17.680519 | 15.676187 | 13.797255 |
| 6 | -16.382671 | 13.917813 | 12.202362 |
| 1 | -16.334267 | 13.662895 | 13.270641 |
| 6 | -15.137231 | 14.754094 | 11.851147 |
| 1 | -15.150578 | 15.673591 | 12.439826 |
| 1 | -15.208488 | 15.057464 | 10.798229 |
| 6 | -16.412038 | 12.61199 | 11.399577 |
| 1 | -17.303161 | 12.020999 | 11.608394 |
| 1 | -16.393031 | 12.816421 | 10.32384 |
| 1 | -15.551115 | 11.988546 | 11.641514 |
| 6 | -13.806332 | 14.031207 | 12.08434 |
| 1 | -13.668215 | 13.190674 | 11.400113 |
| 1 | -12.965054 | 14.714976 | 11.935587 |
| 1 | -13.742008 | 13.640119 | 13.104755 |

Table S5. Interacting structure between L-isoleucine and $(FK)_2$ model

| Atomic Number | Coordinates (Angstroms) | | |
|---|---|---|---|
| | X | Y | Z |
| 6 | -15.259534 | 9.688463 | 9.497417 |
| 6 | -14.774745 | 8.58303 | 10.425168 |
| 6 | -16.548921 | 10.271645 | 10.113229 |
| 8 | -15.294976 | 7.455708 | 10.397469 |
| 6 | -17.168743 | 11.408556 | 9.296686 |
| 6 | -18.412711 | 11.958024 | 9.996266 |
| 6 | -19.187147 | 12.947763 | 9.141827 |
| 7 | -13.795775 | 8.906683 | 11.291902 |
| 6 | -13.314279 | 7.98995 | 12.327884 |
| 6 | -14.229241 | 8.112464 | 13.564849 |
| 6 | -11.844909 | 8.27374 | 12.654886 |
| 8 | -13.918857 | 8.791569 | 14.549017 |
| 6 | -11.246203 | 7.245427 | 13.590551 |
| 6 | -10.848625 | 7.588863 | 14.886625 |
| 6 | -11.082362 | 5.919236 | 13.169279 |
| 6 | -10.301284 | 6.632575 | 15.743381 |
| 6 | -10.539231 | 4.960105 | 14.022766 |
| 6 | -10.146973 | 5.314123 | 15.315118 |
| 7 | -15.39069 | 7.440247 | 13.452552 |
| 6 | -16.473399 | 7.526091 | 14.419811 |
| 6 | -17.150014 | 6.171884 | 14.522515 |
| 6 | -17.53023 | 8.590747 | 14.044902 |
| 8 | -17.090037 | 5.298359 | 13.680707 |
| 6 | -16.978166 | 10.017085 | 14.066272 |
| 6 | -18.066982 | 11.053044 | 13.781476 |
| 6 | -17.518787 | 12.472346 | 13.691305 |

| | | | |
|---|---|---|---|
| 1 | -16.322064 | 10.633854 | 11.12162 |
| 1 | -17.278635 | 9.460819 | 10.221996 |
| 1 | -17.448178 | 11.044865 | 8.302138 |
| 1 | -16.433589 | 12.208479 | 9.150471 |
| 1 | -18.127517 | 12.43667 | 10.936639 |
| 1 | -19.075678 | 11.127419 | 10.245926 |
| 1 | -19.505393 | 12.497194 | 8.201239 |
| 1 | -18.606382 | 13.844111 | 8.922559 |
| 1 | -14.505243 | 10.477792 | 9.450621 |
| 1 | -11.756256 | 9.274014 | 13.084473 |
| 1 | -11.297354 | 8.269997 | 11.708332 |
| 1 | -10.971786 | 8.61085 | 15.227838 |
| 1 | -11.380577 | 5.637438 | 12.164341 |
| 1 | -9.998207 | 6.917965 | 16.744854 |
| 1 | -10.419341 | 3.938607 | 13.678762 |
| 1 | -9.723798 | 4.569662 | 15.980255 |
| 1 | -13.40916 | 6.980806 | 11.92676 |
| 1 | -13.462864 | 9.8624 | 11.313492 |
| 1 | -18.361337 | 8.507622 | 14.750894 |
| 1 | -17.922984 | 8.352658 | 13.051084 |
| 1 | -16.176579 | 10.113796 | 13.327819 |
| 1 | -16.529095 | 10.217752 | 15.045229 |
| 1 | -18.830271 | 11.006347 | 14.56648 |
| 1 | -18.572232 | 10.807261 | 12.841768 |
| 1 | -16.821934 | 12.525766 | 12.842405 |
| 1 | -16.935318 | 12.69916 | 14.589459 |
| 1 | -16.046004 | 7.768763 | 15.393325 |
| 1 | -15.594644 | 6.995617 | 12.563528 |
| 7 | -20.434802 | 13.374264 | 9.859766 |
| 1 | -20.203749 | 13.834008 | 10.743424 |
| 1 | -20.983329 | 14.027976 | 9.298889 |
| 7 | -18.609401 | 13.46169 | 13.589278 |
| 1 | -18.224857 | 14.360706 | 13.31234 |
| 1 | -19.244889 | 13.181065 | 12.844816 |
| 8 | -17.873008 | 6.063224 | 15.648379 |
| 1 | -18.337415 | 5.208512 | 15.649995 |
| 7 | -15.402843 | 9.121948 | 8.148583 |
| 1 | -16.119576 | 8.399682 | 8.177636 |
| 1 | -15.753111 | 9.844386 | 7.526287 |
| 8 | -21.344496 | 11.799946 | 12.592825 |
| 6 | -21.421372 | 10.855963 | 11.764569 |
| 6 | -21.180674 | 9.423922 | 12.282118 |
| 1 | -20.757158 | 9.53102 | 13.288736 |
| 6 | -22.516106 | 8.652319 | 12.41731 |
| 1 | -22.929445 | 8.543674 | 11.406335 |
| 6 | -23.532582 | 9.440029 | 13.266514 |
| 1 | -23.118603 | 9.586339 | 14.272053 |

| | | | |
|---|---|---|---|
| 1 | -23.659115 | 10.439186 | 12.838636 |
| 6 | -24.913353 | 8.784794 | 13.367963 |
| 1 | -25.615499 | 9.446145 | 13.884227 |
| 1 | -24.885367 | 7.84241 | 13.920574 |
| 1 | -25.321904 | 8.577398 | 12.373104 |
| 6 | -22.267763 | 7.252931 | 12.998958 |
| 1 | -21.584825 | 6.664631 | 12.381559 |
| 1 | -21.83975 | 7.326917 | 14.005249 |
| 1 | -23.196975 | 6.683846 | 13.070288 |
| 7 | -20.253002 | 8.710815 | 11.379017 |
| 1 | -19.409529 | 9.26915 | 11.269514 |
| 1 | -19.942909 | 7.861445 | 11.842851 |
| 8 | -21.689913 | 11.013765 | 10.529739 |
| 1 | -21.027998 | 12.53092 | 10.09256 |

Table S6. Interacting structure between D-isoleucine and $(FK)_2$ model

| Atomic Number | Coordinates (Angstroms) | | |
|---|---|---|---|
| | X | Y | Z |
| 8 | 1.923655 | -9.987779 | 9.163181 |
| 1 | 1.969033 | -10.349413 | 10.057295 |
| 8 | 3.30774 | -10.272942 | 6.782691 |
| 1 | 2.498387 | -10.758403 | 6.99496 |
| 8 | 2.442312 | -7.83832 | 5.432136 |
| 1 | 3.291966 | -7.358381 | 5.375191 |
| 8 | 2.943978 | -7.918198 | 9.59583 |
| 6 | 3.239439 | -9.00407 | 7.430055 |
| 1 | 4.211938 | -8.539443 | 7.267796 |
| 6 | 2.165276 | -8.091578 | 6.809293 |
| 1 | 1.199818 | -8.606069 | 6.83405 |
| 6 | 2.051012 | -6.800099 | 7.619666 |
| 6 | 1.853868 | -7.119191 | 9.097489 |
| 1 | 0.90363 | -7.639194 | 9.247291 |
| 1 | 1.849388 | -6.204077 | 9.683566 |
| 6 | 4.696383 | -4.241777 | 5.06459 |
| 6 | 5.202617 | -5.435671 | 5.856831 |
| 6 | 3.202834 | -4.08209 | 5.4328 |
| 8 | 4.966612 | -6.600734 | 5.488107 |
| 6 | 2.580885 | -2.762419 | 4.975963 |
| 6 | 1.06203 | -2.750321 | 5.178851 |
| 6 | 0.441688 | -1.376576 | 4.955979 |
| 7 | 5.830316 | -5.163274 | 7.011589 |
| 6 | 6.02986 | -6.165899 | 8.055044 |
| 6 | 5.929723 | -5.438902 | 9.395911 |
| 6 | 7.378487 | -6.911826 | 7.932761 |
| 8 | 6.608395 | -4.421937 | 9.602861 |
| 6 | 7.488871 | -7.990536 | 8.984016 |

| | | | |
|---|---|---|---|
| 6 | 8.183129 | -7.762786 | 10.177312 |
| 6 | 6.813256 | -9.206209 | 8.82086 |
| 6 | 8.197022 | -8.723822 | 11.188533 |
| 6 | 6.824074 | -10.168429 | 9.829386 |
| 6 | 7.513939 | -9.928223 | 11.019054 |
| 7 | 5.084942 | -5.978207 | 10.284372 |
| 6 | 4.754645 | -5.330614 | 11.541198 |
| 6 | 4.842745 | -6.336645 | 12.673952 |
| 6 | 3.349133 | -4.678554 | 11.497472 |
| 8 | 4.976757 | -7.535296 | 12.540025 |
| 6 | 3.177211 | -3.750351 | 10.293012 |
| 6 | 1.759101 | -3.192759 | 10.155847 |
| 6 | 1.633526 | -2.274281 | 8.943806 |
| 1 | 3.091908 | -4.157347 | 6.520096 |
| 1 | 2.658861 | -4.928585 | 5.000741 |
| 1 | 2.801369 | -2.579394 | 3.917889 |
| 1 | 3.038529 | -1.940102 | 5.539214 |
| 1 | 0.824082 | -3.080811 | 6.196949 |
| 1 | 0.598637 | -3.475428 | 4.500048 |
| 1 | 0.657064 | -1.03591 | 3.938588 |
| 1 | 0.919759 | -0.658953 | 5.638864 |
| 1 | 5.236401 | -3.345258 | 5.379409 |
| 1 | 8.191801 | -6.188813 | 8.027649 |
| 1 | 7.418852 | -7.342759 | 6.929905 |
| 1 | 8.701755 | -6.820261 | 10.31961 |
| 1 | 6.268577 | -9.394184 | 7.901746 |
| 1 | 8.734895 | -8.528792 | 12.10981 |
| 1 | 6.291516 | -11.102469 | 9.688002 |
| 1 | 7.518721 | -10.673458 | 11.806659 |
| 1 | 5.22322 | -6.89066 | 7.970091 |
| 1 | 6.013641 | -4.197421 | 7.252431 |
| 1 | 3.203406 | -4.125092 | 12.42789 |
| 1 | 2.595628 | -5.472167 | 11.474193 |
| 1 | 3.419346 | -4.294614 | 9.37499 |
| 1 | 3.896548 | -2.926476 | 10.365947 |
| 1 | 1.47212 | -2.648624 | 11.062798 |
| 1 | 1.052317 | -4.025784 | 10.047828 |
| 1 | 2.042249 | -2.792734 | 8.065379 |
| 1 | 2.250737 | -1.383897 | 9.095764 |
| 1 | 5.501016 | -4.557151 | 11.727334 |
| 1 | 4.490371 | -6.75373 | 9.991237 |
| 7 | -1.023814 | -1.418563 | 5.123862 |
| 1 | -1.232634 | -1.674285 | 6.086823 |
| 1 | -1.401733 | -0.483023 | 4.999464 |
| 7 | 0.241819 | -1.834995 | 8.733509 |
| 1 | 0.212242 | -1.216625 | 7.926291 |
| 1 | -0.314077 | -2.644949 | 8.465018 |

| | | | |
|---|---|---|---|
| 8 | 4.732008 | -5.735866 | 13.871543 |
| 1 | 4.76241 | -6.409503 | 14.572689 |
| 7 | 4.964285 | -4.457266 | 3.636602 |
| 1 | 4.457004 | -5.288379 | 3.339011 |
| 1 | 4.567344 | -3.682737 | 3.111702 |
| 6 | 3.062847 | -9.178544 | 8.946271 |
| 1 | 3.951642 | -9.639419 | 9.379451 |
| 8 | 0.933437 | -6.045079 | 7.138907 |
| 1 | 1.099214 | -5.110578 | 7.317267 |
| 1 | 2.973291 | -6.224695 | 7.496477 |

Table S7. Interacting structure between L-xylose and $(FK)_2$ model

| Atomic Number | Coordinates (Angstroms) | | |
|---|---|---|---|
| | X | Y | Z |
| 8 | 3.000962 | -8.289455 | 8.100275 |
| 1 | 2.235477 | -7.701223 | 8.065524 |
| 8 | 3.738145 | -10.857467 | 8.976862 |
| 1 | 3.920463 | -10.083909 | 9.537514 |
| 8 | 6.096833 | -11.141127 | 7.364354 |
| 1 | 6.996196 | -10.789352 | 7.412052 |
| 8 | 2.799263 | -9.066164 | 5.896575 |
| 6 | 3.803709 | -10.483666 | 7.605561 |
| 1 | 3.53775 | -11.377309 | 7.035867 |
| 6 | 5.203169 | -10.038235 | 7.175933 |
| 1 | 5.524165 | -9.207415 | 7.809184 |
| 6 | 5.181591 | -9.585234 | 5.71024 |
| 6 | 4.078124 | -8.552765 | 5.48075 |
| 1 | 4.304405 | -7.632148 | 6.023634 |
| 1 | 3.985306 | -8.320369 | 4.420668 |
| 6 | 5.226968 | -4.175101 | 5.4985 |
| 6 | 6.04865 | -5.306575 | 6.096 |
| 6 | 3.743212 | -4.603632 | 5.607852 |
| 8 | 6.444876 | -6.255871 | 5.400963 |
| 6 | 2.758035 | -3.522983 | 5.160146 |
| 6 | 1.30379 | -3.951968 | 5.371676 |
| 6 | 0.305153 | -2.879724 | 4.951864 |
| 7 | 6.238085 | -5.257233 | 7.427173 |
| 6 | 6.656076 | -6.438008 | 8.189802 |
| 6 | 6.062033 | -6.263518 | 9.592819 |
| 6 | 8.186699 | -6.619086 | 8.22094 |
| 8 | 6.564304 | -5.456363 | 10.385375 |
| 6 | 8.572165 | -8.015493 | 8.653341 |
| 6 | 8.502357 | -8.406619 | 9.996424 |
| 6 | 8.958025 | -8.96575 | 7.700293 |
| 6 | 8.797006 | -9.716524 | 10.375263 |
| 6 | 9.265297 | -10.274315 | 8.076694 |

| | | | |
|---|---|---|---|
| 6 | 9.17772 | -10.655891 | 9.416637 |
| 7 | 4.967186 | -6.994183 | 9.857251 |
| 6 | 4.143874 | -6.761015 | 11.031381 |
| 6 | 3.84536 | -8.074732 | 11.727978 |
| 6 | 2.831152 | -6.019144 | 10.680525 |
| 8 | 4.051098 | -9.181742 | 11.270386 |
| 6 | 3.081662 | -4.691026 | 9.963626 |
| 6 | 1.781486 | -3.972893 | 9.596156 |
| 6 | 2.035028 | -2.654345 | 8.874088 |
| 1 | 3.530708 | -4.866354 | 6.650514 |
| 1 | 3.598945 | -5.512473 | 5.01214 |
| 1 | 2.912728 | -3.285497 | 4.101467 |
| 1 | 2.953395 | -2.601132 | 5.721261 |
| 1 | 1.148813 | -4.191411 | 6.431473 |
| 1 | 1.105263 | -4.872405 | 4.809972 |
| 1 | 0.403921 | -2.687839 | 3.879276 |
| 1 | 0.554907 | -1.940989 | 5.466719 |
| 1 | 5.369026 | -3.274868 | 6.102849 |
| 1 | 8.616957 | -5.865742 | 8.884938 |
| 1 | 8.5585 | -6.430759 | 7.21287 |
| 1 | 8.214672 | -7.682496 | 10.751145 |
| 1 | 9.014898 | -8.678725 | 6.656163 |
| 1 | 8.73204 | -10.001715 | 11.419503 |
| 1 | 9.569744 | -10.993901 | 7.324325 |
| 1 | 9.407765 | -11.673865 | 9.710038 |
| 1 | 6.212727 | -7.304871 | 7.703214 |
| 1 | 5.858684 | -4.467232 | 7.933241 |
| 1 | 2.286063 | -5.84436 | 11.611318 |
| 1 | 2.209091 | -6.670751 | 10.059563 |
| 1 | 3.659851 | -4.866232 | 9.050623 |
| 1 | 3.693604 | -4.044458 | 10.60316 |
| 1 | 1.189475 | -3.786012 | 10.499565 |
| 1 | 1.177413 | -4.623275 | 8.950048 |
| 1 | 2.662469 | -2.848591 | 7.993861 |
| 1 | 2.604844 | -1.985801 | 9.526606 |
| 1 | 4.71887 | -6.149684 | 11.728063 |
| 1 | 4.545214 | -7.545493 | 9.112752 |
| 7 | -1.08225 | -3.309423 | 5.213423 |
| 1 | -1.205301 | -3.418763 | 6.218016 |
| 1 | -1.719657 | -2.567156 | 4.937114 |
| 7 | 0.769353 | -1.982228 | 8.52331 |
| 1 | 0.978505 | -1.142148 | 7.989722 |
| 1 | 0.25547 | -2.582166 | 7.881785 |
| 8 | 3.299975 | -7.875507 | 12.936571 |
| 1 | 3.094948 | -8.735663 | 13.341808 |
| 7 | 5.70356 | -3.89257 | 4.140708 |
| 1 | 5.583543 | -4.731116 | 3.576383 |

| | | | |
|---|---|---|---|
| 1 | 5.106253 | -3.183239 | 3.726084 |
| 6 | 2.75691 | -9.410712 | 7.271396 |
| 1 | 1.751398 | -9.798802 | 7.442982 |
| 1 | 4.984984 | -10.467096 | 5.09349 |
| 8 | 6.449517 | -9.075845 | 5.292217 |
| 1 | 6.517656 | -8.135167 | 5.543715 |

Table S8. Interacting structure between D-xylose and $FK_2$ model

#### 5. NMR Spectroscopy

### 5.1 $(FK)_4$

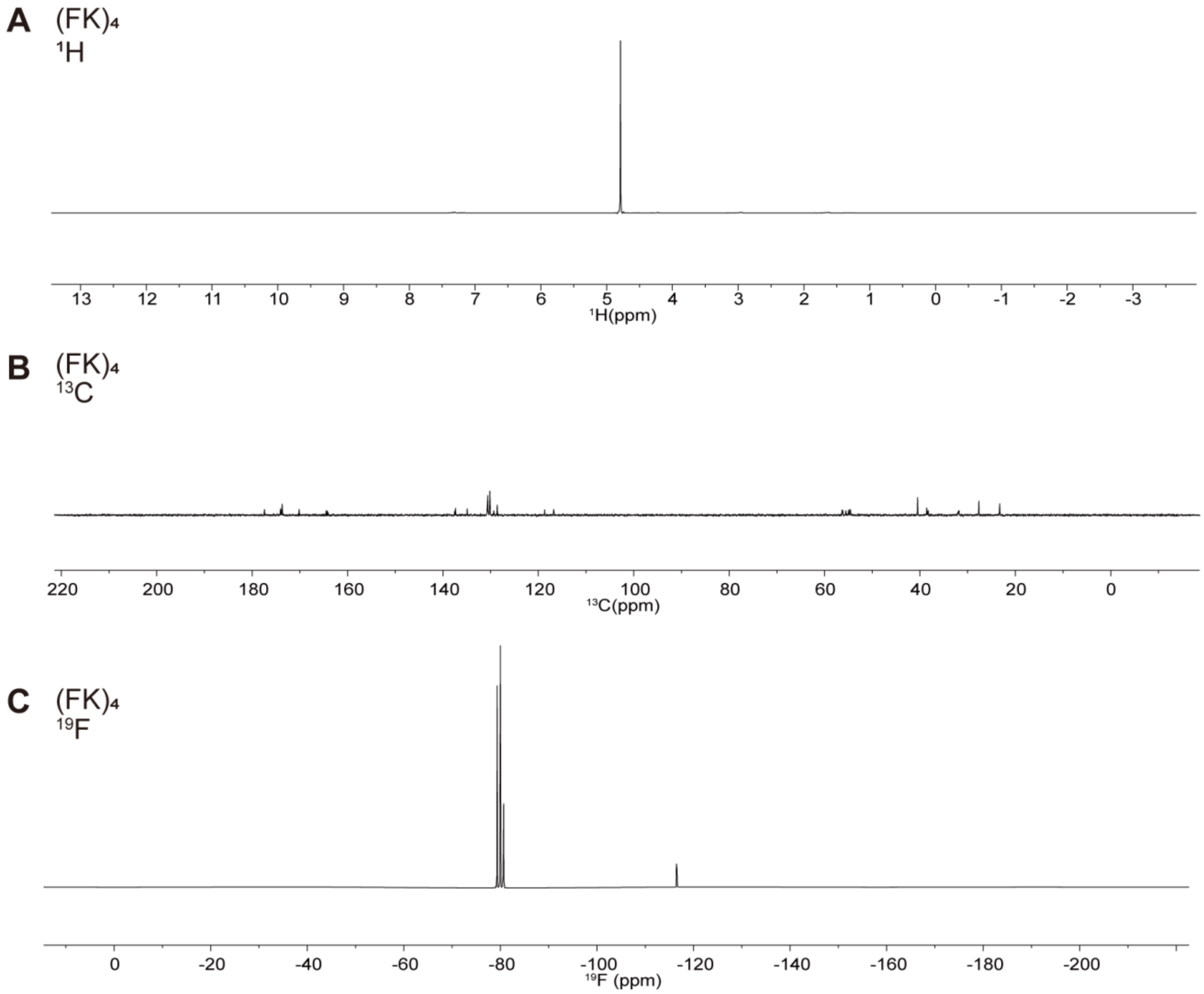

Figure S6. NMR of $(FK)_4$. A. $^{1}H$ NMR spectroscopy with presence of 140 mg/ml $(FK)_4$ in $D_2O$. B. Proton decoupled $^{13}C$ NMR spectroscopy with 140 mg/ml $(FK)_4$. C. $^{19}F$ NMR spectroscopy with 140 mg/ml $(FK)_4$.

#### 5.2 Proline

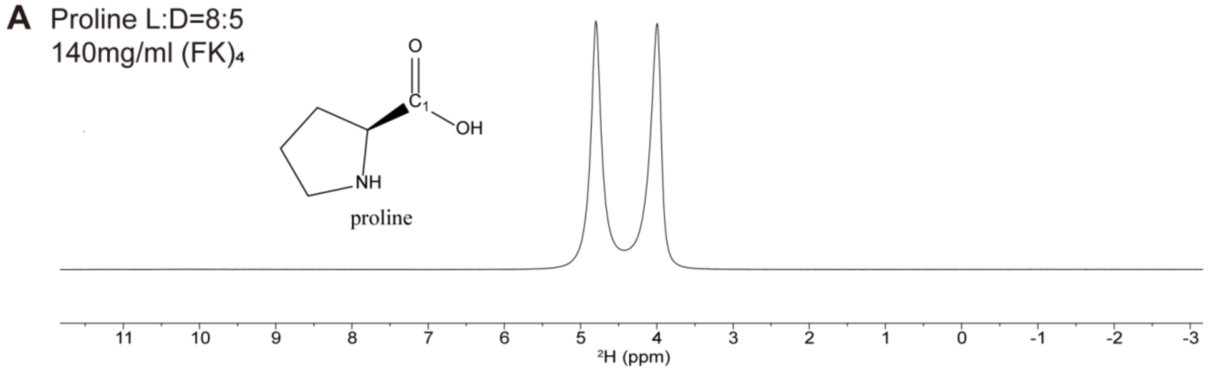

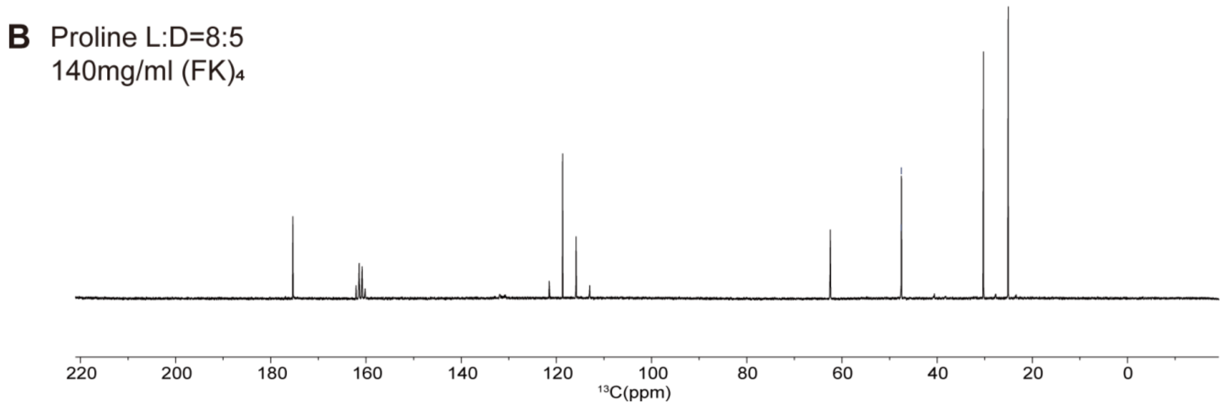

Figure S7. NMR of proline with $(FK)_4$. A. $^2H$ NMR spectroscopy with presence of 140 mg/ml $(FK)_4$ in $D_2O$. B. Proton decoupled $^{13}C$ NMR spectroscopy of 65 mM proline enantiomers with 140 mg/ml $(FK)_4$.

#### 5.3 Valine

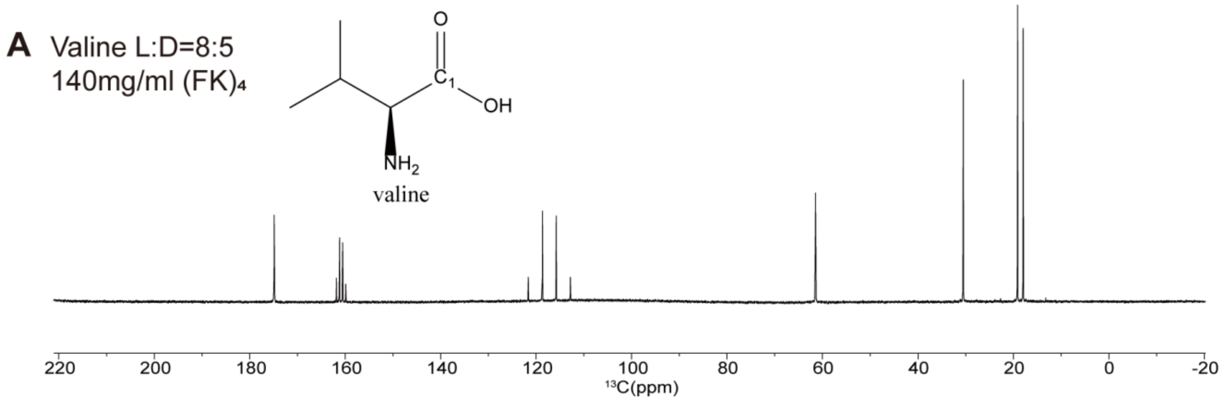

Figure S8. NMR of valine with $(FK)_4$. A. Proton decoupled $^{13}C$ NMR spectroscopy of 65 mM valine enantiomers with 140 mg/ml $(FK)_4$ in $D_2O$.

#### 5.4 Leucine

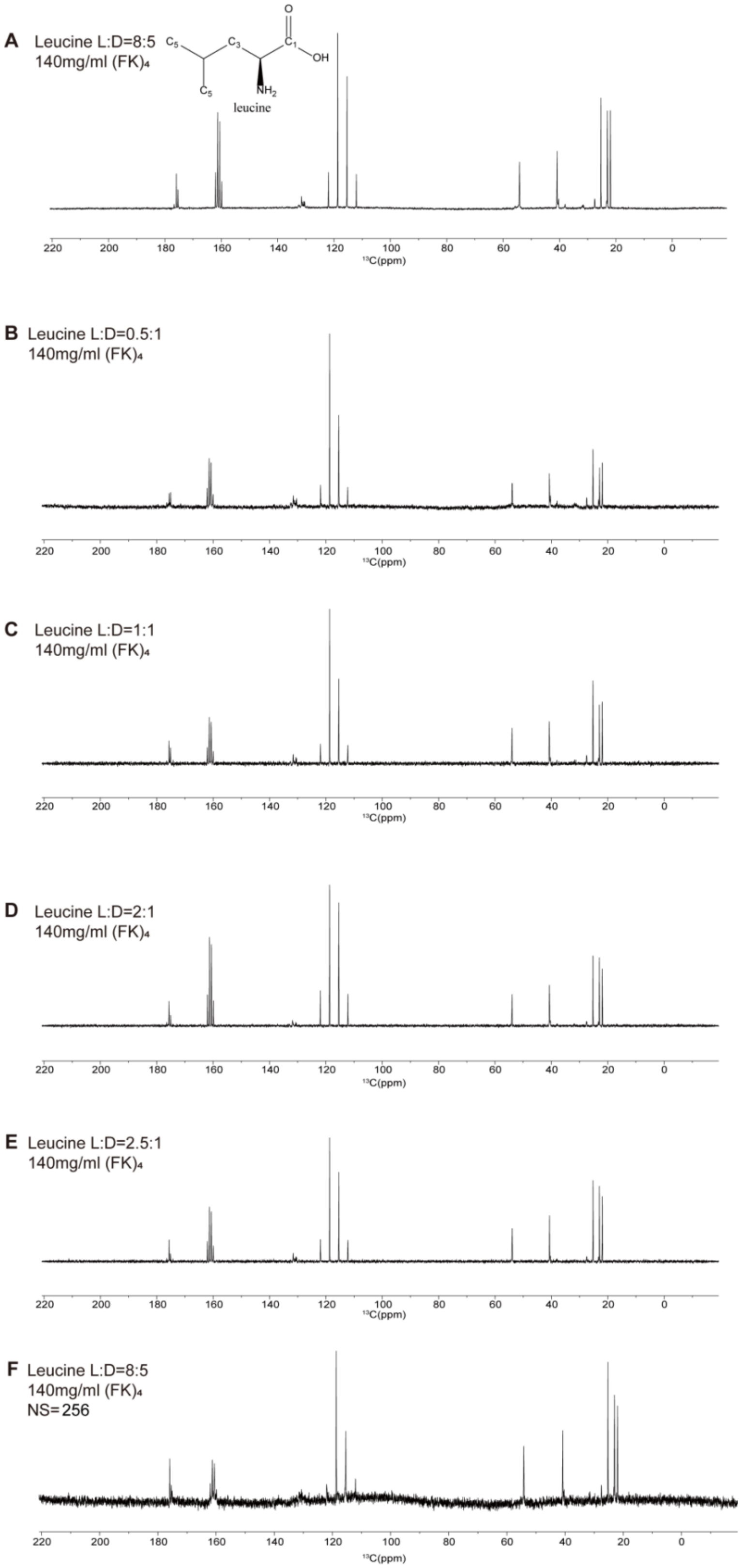

Figure S9. Proton decoupled $^{13}$C NMR spectroscopy of 65 mM leucine enantiomers with 140 mg/ml $(FK)_4$ in $D_2O$. A-E. The molar ratio of the L- and D-enantiomers are 8:5, 0.5:1, 1:1, 2:1 and 2.5:1. F. Superimposition of 256 number of scans.

#### 5.5 Isoleucine

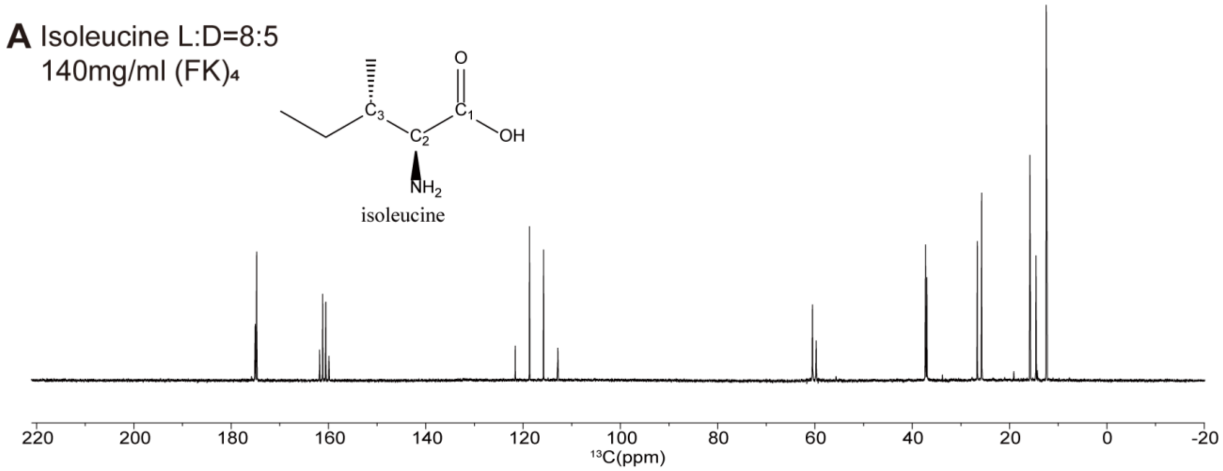

Figure S10. NMR of isoleucine with $(FK)_4$. A. Proton decoupled $^{13}C$ NMR spectroscopy of 65 mM isoleucine enantiomers with 140 mg/ml $(FK)_4$ in $D_2O$.

#### 5.6 Phenylalanine

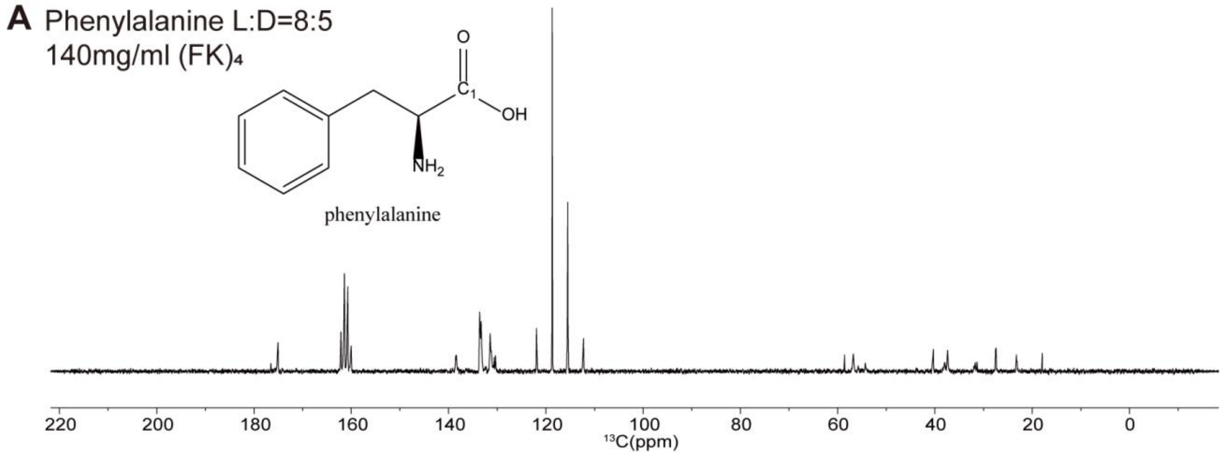

Figure S11. NMR of phenylalanine with $(FK)_4$. A. Proton decoupled $^{13}C$ NMR spectroscopy of 65 mM phenylalanine enantiomers with 140 mg/ml $(FK)_4$ in $D_2O$.

#### 5.7 Glutamine

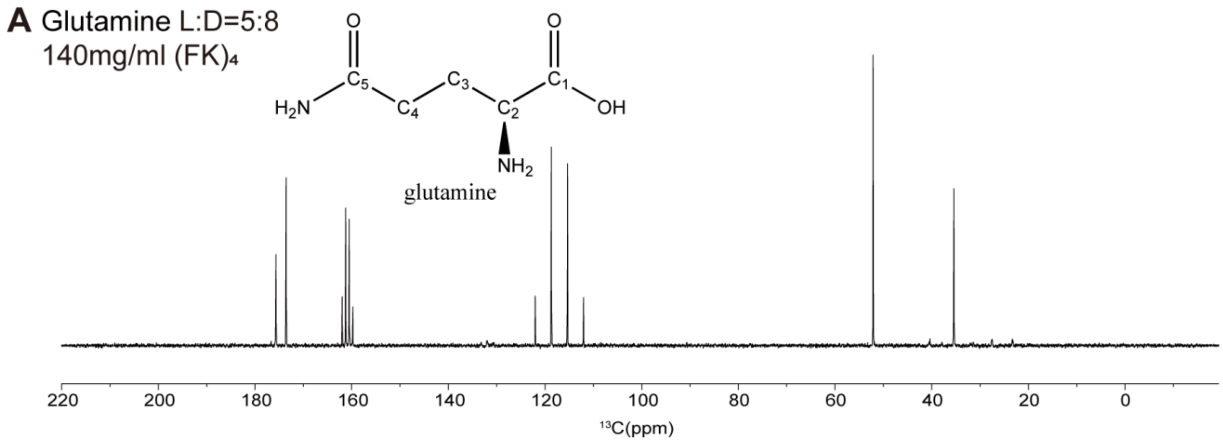

Figure S12. NMR of glutamine with $(FK)_4$. A. Proton decoupled $^{13}C$ NMR spectroscopy of 65 mM glutamine enantiomers with 140 mg/ml $(FK)_4$ in $D_2O$.

#### 5.8 Glutamic acid

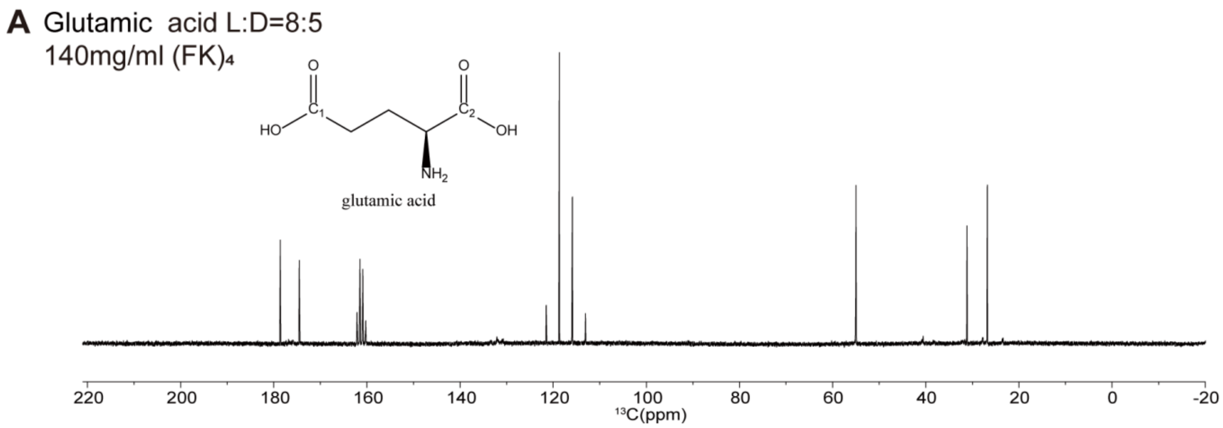

Figure S13. NMR of glutamic acid with $(FK)_4$. A. Proton decoupled $^{13}C$ NMR spectroscopy of 65 mM glutamic acid enantiomers with 140 mg/ml $(FK)_4$ in $D_2O$.

#### 5.9 Aspartic acid

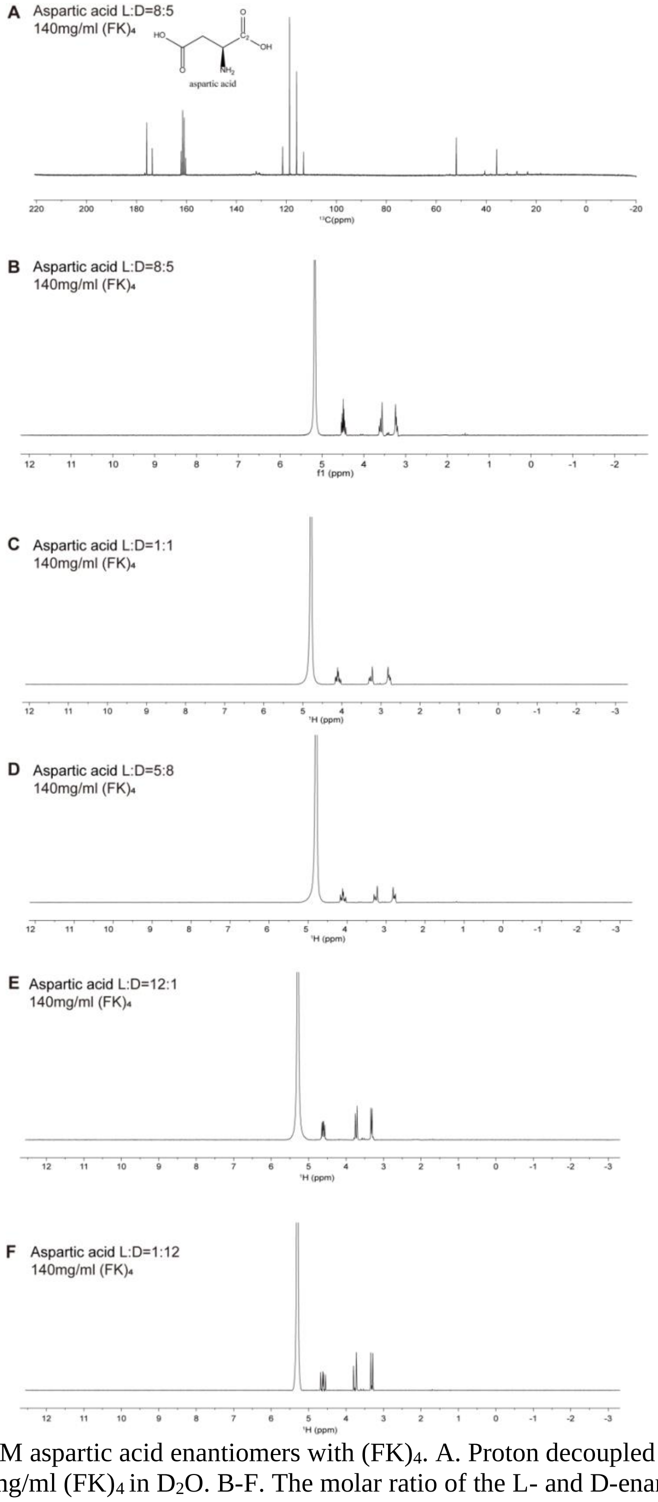

Figure S14. NMR of 65 mM aspartic acid enantiomers with $(FK)_4$. A. Proton decoupled $^{13}C$ NMR spectroscopy of aspartic acid with 140 mg/ml $(FK)_4$ in $D_2O$. B-F. The molar ratio of the L- and D-enantiomers are 8:5, 1:1, 5:8, 12:1 and 1:12.

#### 5.10 Glucose

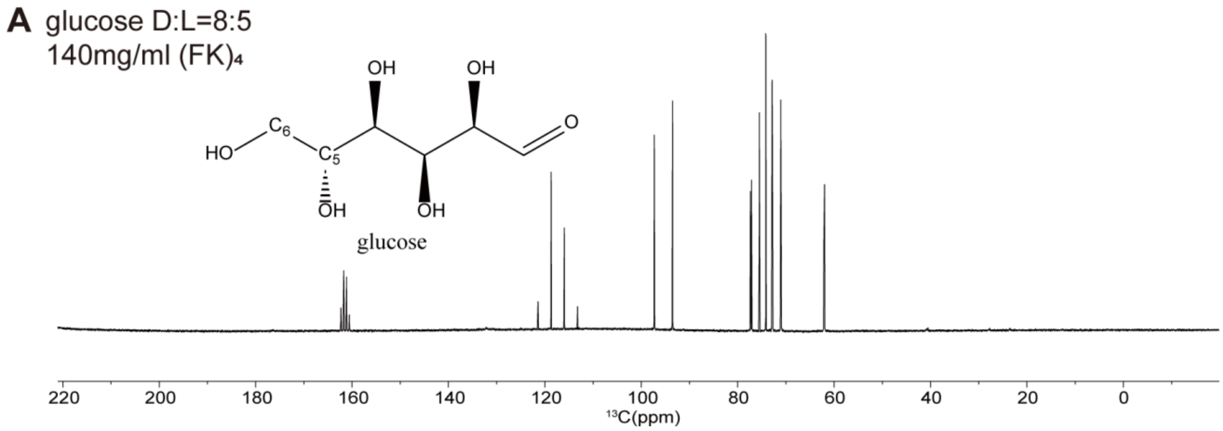

Figure S15. NMR of glucose with $(FK)_4$. A. Proton decoupled $^{13}C$ NMR spectroscopy of 65 mM glucose enantiomers with 140 mg/ml $(FK)_4$ in $D_2O$.

#### 5.11 Frutose

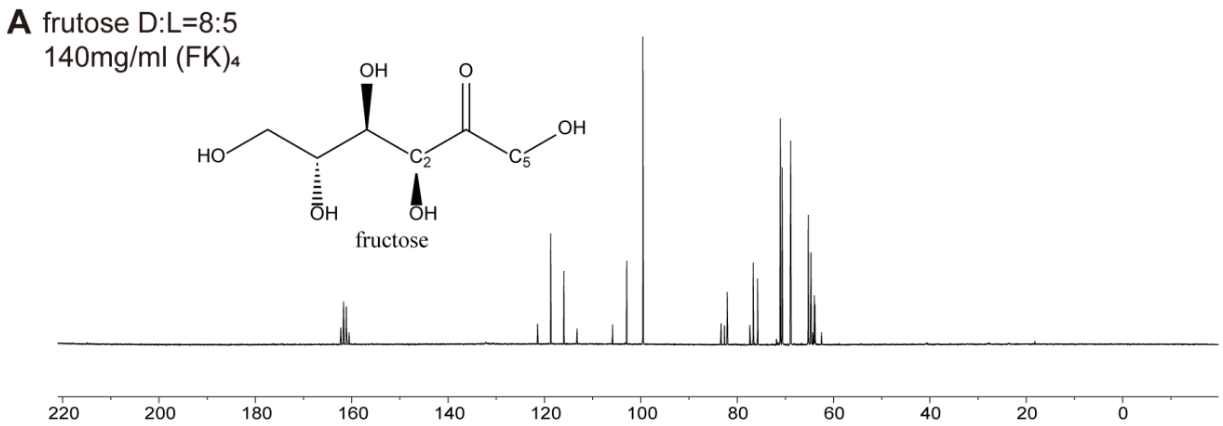

Figure S16. NMR of frutose with $(FK)_4$. A. Proton decoupled $^{13}C$ NMR spectroscopy of 65 mM fructose enantiomers with 140 mg/ml $(FK)_4$ in $D_2O$.

#### 5.12 Galactose

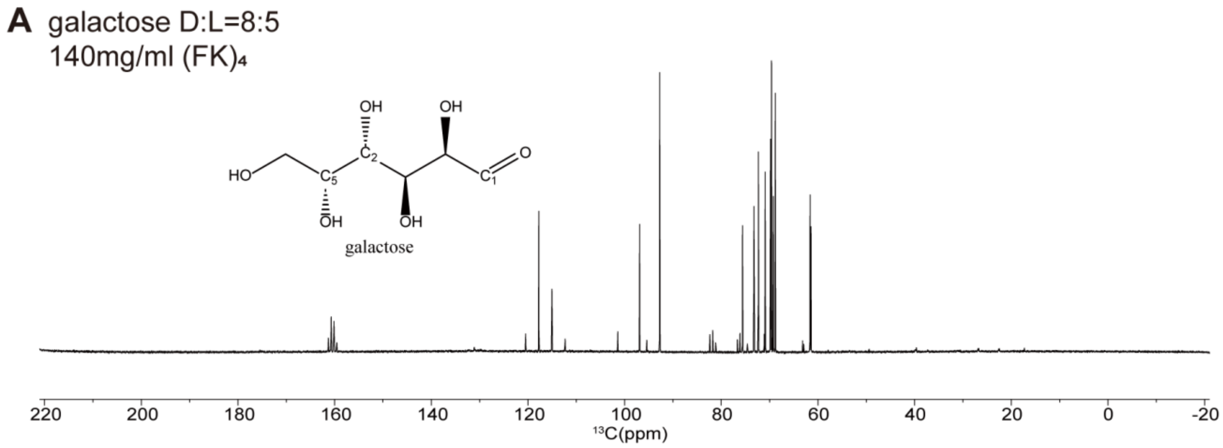

Figure S17. NMR of galactose with $(FK)_4$. A. Proton decoupled $^{13}C$ NMR spectroscopy of 65 mM galactose enantiomers with 140 mg/ml $(FK)_4$ in $D_2O$.

#### 5.13 Xylose

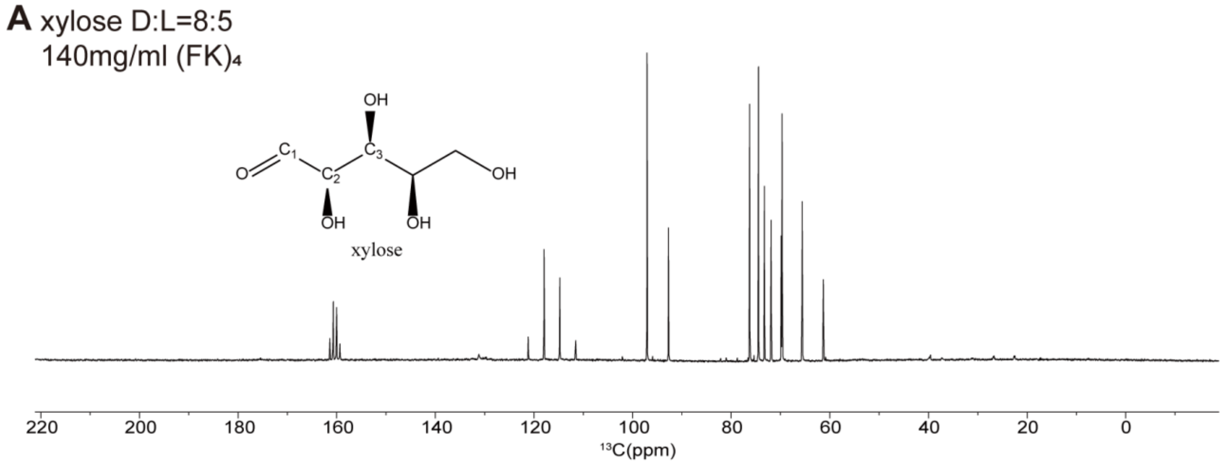

Figure S18. NMR of xylose with $(FK)_4$. A. Proton decoupled $^{13}C$ NMR spectroscopy of 65 mM xylose enantiomers with 140 mg/ml $(FK)_4$ in $D_2O$.

#### 5.14 Threitol

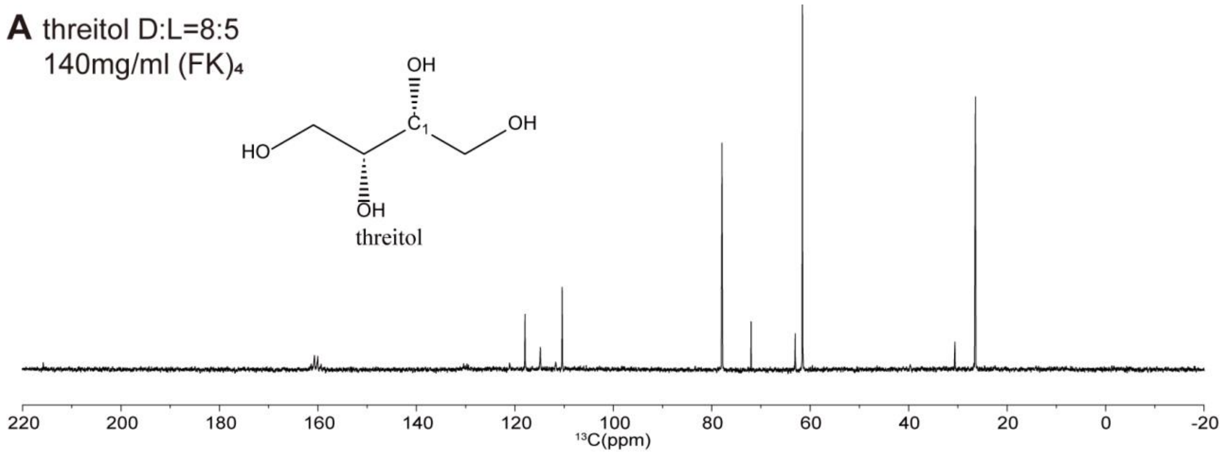

Figure S19. NMR of threitol with $(FK)_4$. A. Proton decoupled $^{13}C$ NMR spectroscopy of 65 mM threitol enantiomers with 140 mg/ml $(FK)_4$ in $D_2O$.

#### 5.15 Lactose

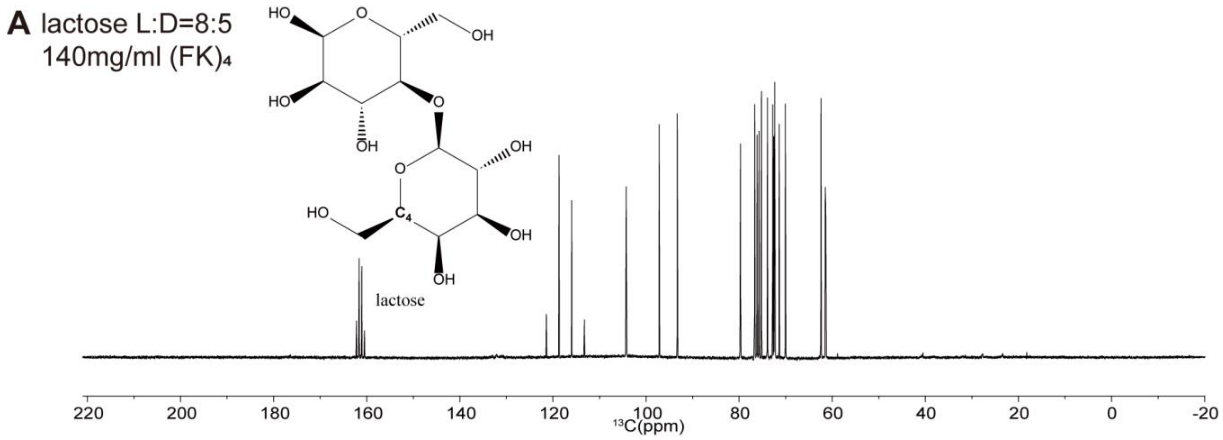

Figure S20. NMR of lactose with $(FK)_4$. A. Proton decoupled $^{13}C$ NMR spectroscopy of 65 mM lactose enantiomers with 140 mg/ml $(FK)_4$ in $D_2O$.

##### 5.16 Lactic acid

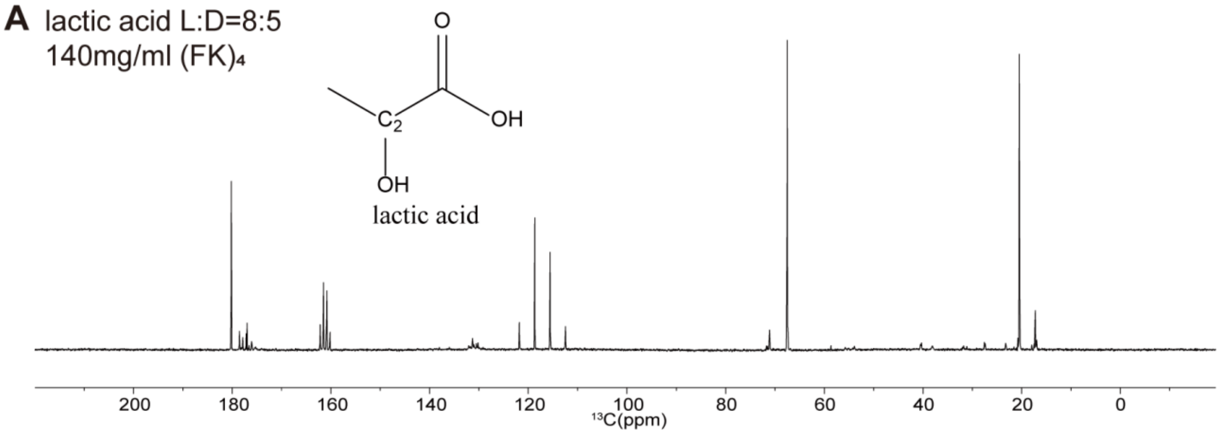

Figure S21. NMR of lactic acid with $(FK)_4$. A. Proton decoupled $^{13}C$ NMR spectroscopy of 65 mM lactic acid enantiomers with 140 mg/ml $(FK)_4$ in $D_2O$.

##### 5.17 2-methylvaleric acid

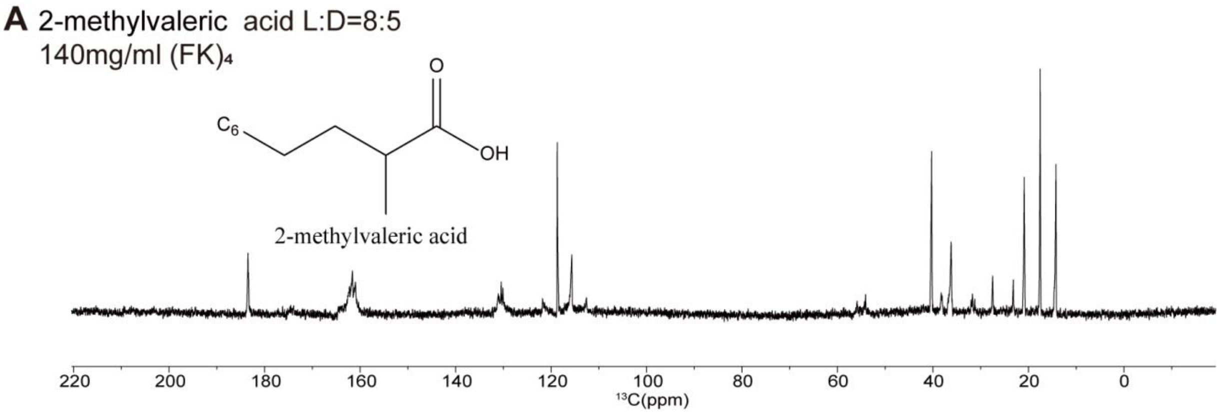

Figure S22. NMR of 2-methylvaleric acid with $(FK)_4$. A. Proton decoupled $^{13}C$ NMR spectroscopy of 65 mM 2-methylvaleric acid enantiomers with 140 mg/ml $(FK)_4$ in $D_2O$.

#### 5.18 Terbutaline

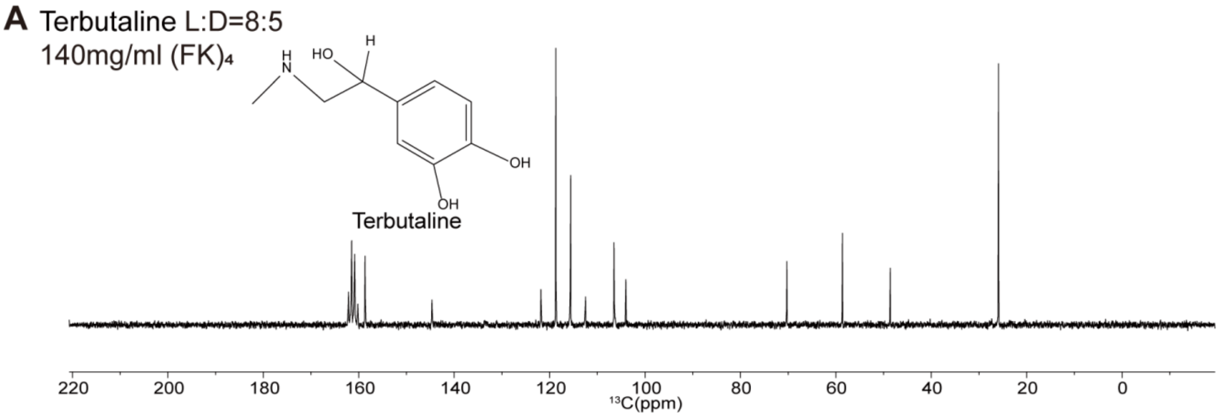

Figure S23. NMR of terbutaline with $(FK)_4$. A. Proton decoupled $^{13}$C NMR spectroscopy of 65 mM terbutaline enantiomers with 140 mg/ml $(FK)_4$ in $D_2O$.

#### 5.19 Epinephrine

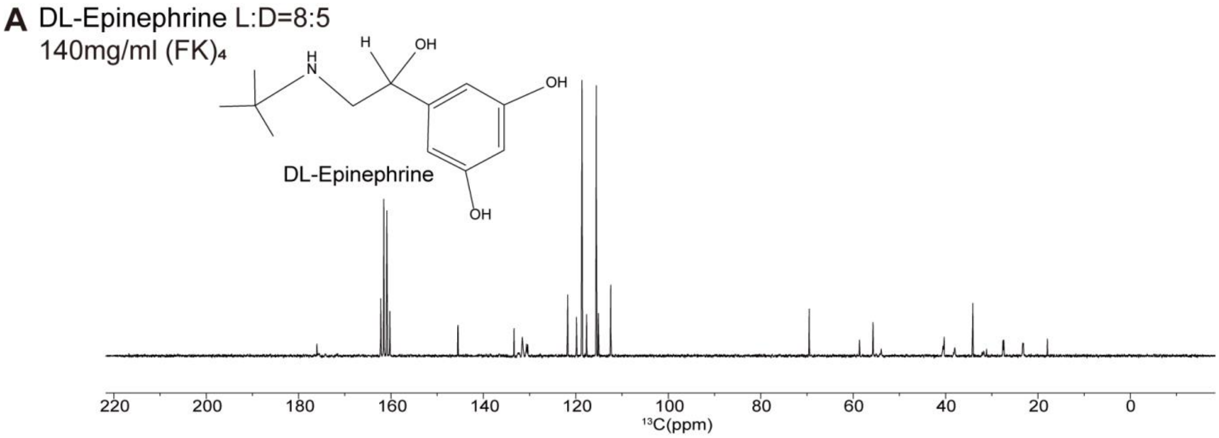

Figure S24. NMR of epinephrine with $(FK)_4$. A. Proton decoupled $^{13}$C NMR spectroscopy of 65 mM epinephrine enantiomers with 140 mg/ml $(FK)_4$ in $D_2O$.

##### 5.20 Chloroquine

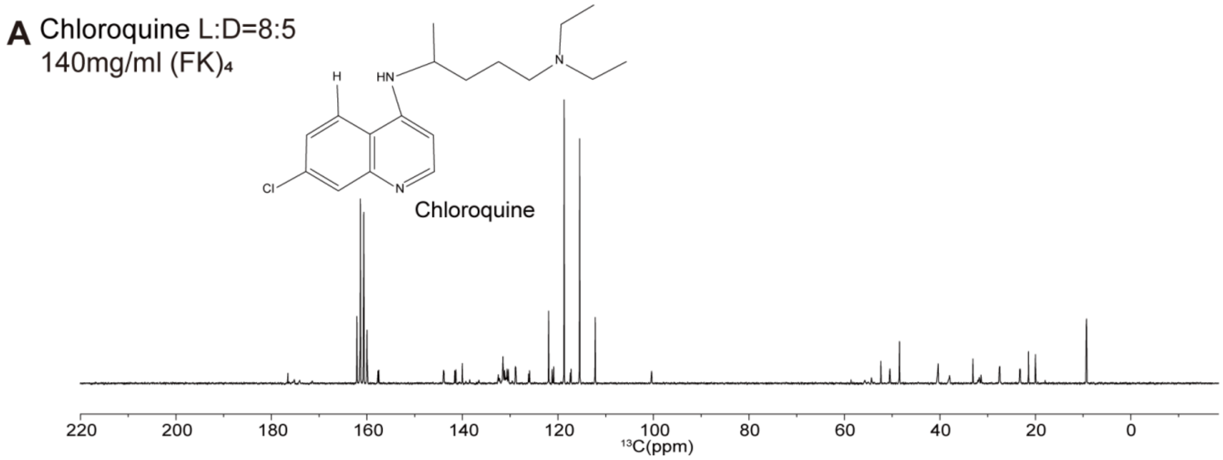

Figure S25. NMR of chloroquine with $(FK)_4$. A. Proton decoupled $^{13}C$ NMR spectroscopy of 65 mM chloroquine enantiomers with 140 mg/ml $(FK)_4$ in $D_2O$.
